# Novel verbal instructions recruit abstract neural patterns of time-variable information dimensionality

**DOI:** 10.1101/2024.07.19.603493

**Authors:** Paula Pena, Ana F. Palenciano, Carlos González-García, María Ruz

**Affiliations:** Mind, Brain and Behavior Research Center, University of Granada, Granada, Spain

**Keywords:** cognitive control, novel instructed behavior, task preparation and implementation, neural geometry

## Abstract

Human performance is endowed by neural representations of information that is relevant for behavior, some of which are also activated in a preparatory fashion to optimize later execution. Most studies to date have focused on highly practiced actions, leaving largely unaddressed the novel re-configuration of information to generate unique whole task-sets. Using electroencephalography (EEG), this study investigated the dynamics of the content and geometry reflected on the neural patterns of control representations during re-configuration of information. We designed a verbal instruction paradigm where each trial involved novel combinations of multi-component task information. By manipulating three task-relevant factors in a sample of 40 participants (26 females, 14 males), we observed complex coding schemes throughout the trial, during both preparation and implementation stages. The temporal profiles were consistent with a hierarchical structure: whereas task information was active in a sustained manner, the coding of more concrete stimulus features was more transient. Data showed both high dimensionality and abstraction, particularly during instruction encoding and target processing. Our results suggest that whenever task content could be recovered from neural patterns of activity, there was evidence of abstract coding, with an underlying geometry that favored generalization. During target processing, where potential interference across stimulus and response factors increased, orthogonal configurations also appeared. Overall, our findings uncover the dynamic manner with which control representations operate during novel recombination unique scenarios, with changes in dimensionality and abstraction adjusting along processing stages.

**Significance Statement:** The neural mechanisms that support task performance in novel contexts have been largely overlooked. Cognitive control is thought to enable complex behavior through the active maintenance of task sets, containing essential information for execution. However, how novel whole combinations of information are organized in neural patterns and their temporal dependencies remain unknown. Here, using a novel complex instruction paradigm, we observed that coding of informational content and its underlying geometry followed a dynamic temporal pattern. Our results reveal varying dimensionality and abstraction throughout the trial, with neural codes generally structured in a geometry favoring generalization of relevant information across task demands. These findings provide a first glimpse into the temporal computations engaged by the brain when encountering novel recombination settings.

## 1. Introduction

Humans excel at following instructed commands, often guided by sentences conveying details about the relevance of information and the rules associating them with the required actions. This ability, linked to cognitive control mechanisms, is most useful in novel and changing scenarios. However, we know very little about how the human brain encodes and recombines information to achieve such success. Theoretical models propose that top-down control organizes and maintains the relevant information by creating “task sets” (Miller & Cohen, 2001; Sakai, 2008), with empirical evidence linking these to the Multiple Demand Network (Duncan, 2010). Recently, the study of the neural geometry of activity patterns in brain regions has gained relevance, examining whether control codes are structured to favor separability or generalization of information (Fusi et al., 2016). Still, the neural mechanisms supporting the flexible generation of novel combinatorial scenarios remain an elusive topic.

Control-related information is anticipated when preparing to implement a task and later reactivated during task implementation (Grootswagers et al., 2018; Sakai & Passingham, 2003). Nevertheless, most of the evidence comes from repetitive and highly practiced tasks, which limits flexibility and constrains extrapolation to more novel contexts, cornerstone of human cognitive control. The small body of research incorporating novel tasks has demonstrated that instruction-related information is coded during both preparation and task implementation (Muhle-Karbe et al., 2017; González-García, 2017; Hartstra et al., 2011; Palenciano et al., 2019a). However, these fMRI studies are blind to the underlying temporal dynamics. Few studies have used electroencephalography (EEG) to investigate instructed behavior (Formica et al., 2021; Formica et al., 2022) but none has employed temporally resolved multivariate analyses to characterize the coding of novel instructions in time.

Characterizing control representations extends to the examination of their underlying configuration, or neural geometry (Badre et al., 2021), which likely contributes to the multifaceted nature of cognitive control. Studies indicate that pattern geometry adapts to task demands, with spatial arrangements of neural activity shifting based on task requirements (Musslick & Cohen, 2021; Kadohisa et al., 2023; Flesch et al., 2022). When reuse of information is needed, control representations may use abstraction, i.e., compressed low-dimensional geometries, to generalize task-relevant components across contexts (Cole et al., 2013; Badre et al., 2021; Verbeke & Verguts, 2022; Bhandari et al., 2024). Conversely, the flexibility of input encoding appears to be supported by a different operation, dimensionality expansion, which improves information separability by maximizing the richness of attributes represented (Rigotti et al., 2013; Fusi et al., 2016). Recent work suggests that representational spaces are initially high-dimensional during novel tasks and decrease with learning (Bhandari et al., 2024; Wojcik et al., 2023; Farrel et al., 2022). Dimensionality further varies at faster timescales, expanding during task execution (Kikumoto et al., 2024a; Bhandari et al., 2024). Moreover, studies examining more detailed geometrical models based on lower-dimensional manifolds have related parallel coding dimensions to better generalization, and orthogonal arrangements with conflict minimization (Stokes et al., 2020; Muhle-Karbe et al., 2023). However, how these geometrical features dynamically adapt during the assembly and execution of novel tasks combinations remains uncertain.

The current study employed verbal instructions that generated trial-by-trial novelty by re-using task components (Cole et al., 2013; Palenciano et al., 2019a), which we expected would favor abstract, multidimensional coding. Using EEG data, we investigated how neural pattern geometry unfolds across trial stages with different requirements, from building whole task sets based on the instructions to applying response contingencies to specific target combinations. The instructions were designed to manipulate task content thought to engage different cognitive processes: task demands (information selection or integration), target category (animate or inanimate), and target relevant feature to respond (color or shape). This enabled analyzing whether abstraction levels differentially varied with task components. We studied dimensionality dynamics across time, comparing preparation and implementation phases. Finally, we explored the spatial arrangement of our task components when constrained to a low-dimensional space, expecting a parallel alignment to support information sharing across contexts.

## 2. Materials and Methods

### 2.1 Participants

Forty participants (mean age = 21.85, range: 18-27, SD = 2.28; 26 females and 14 males) received economic compensation for taking part in the experiment, varying from 30 to 35 euros, according to their average performance on the task. They were all native Spanish speakers, right-handed, with normal or corrected vision and no history of neurological issues. Participants that were not able to complete the practice session to criterion (see below) received a compensation of 5 euros. Additionally, the data of one participant had to be excluded due to low quality of the EEG recording, resulting in a final sample size of 39. All participants signed a consent form approved by the local Ethics Committee (reference 1584/CEIH/2020). The experiment was carried out at the Mind, Brain and Behavior Research Center (CIMCYC) of the University of Granada.

The sample size was selected based on a previous fMRI study where a similar experimental paradigm was employed (Palenciano et al., in preparation). We compared our targeted sample size against the one obtained with a power analysis focused on a behavioral effect size. This choice was due to the difficulty of finding reliable effect sizes from previous studies employing similar multivariate approaches. We used the software PANGEA (Westfall, 2015) to detect a modest effect size (Cohen’s d = 0.3) of the variable Task Demand in the behavioral data (Reaction Times and Accuracy). Our sample size of 40 would achieve a statistical power of 90,5%.

### 2.2 Experimental Design

The experiment consisted of a main instruction-following task with a 2×2×2 within-subject design, where the independent variables were manipulated according to different hierarchical levels, with Task Demand (integration vs. selection) setting the goal of the task, and Target Category (animate vs. inanimate) and Target Relevant Feature (color vs. shape) as lower-level variables. The dependent variables were behavioral (Reaction Times and Accuracy) and electrophysiological (Voltage Values). Additionally, we ran a localizer task with a 4×2 within-subject design, manipulating the Target Subcategory of the stimuli (sea animal, land animal, musical instrument or tool) and Trial Type (repeat or non-repeat trials). This task was implemented for other analyses, not included in the current article, and thus will no further be discussed.

### 2.3 Apparatus and Stimuli

We generated 512 verbal instructions, which were novel as a whole, by recombining the elements derived from the three independent variables and the instructed response. Each instruction consisted of an “if… then” statement, indicating a condition about the two upcoming targets, together with the required response in case the condition was fulfilled. After the instructions, participants were shown pairs of targets surrounded by frames, which were generated by combining 8 images, 4 colors and 4 shapes. Images were taken from a total pool of 16 and were either animate (land animals: horse, bear, dog or cat and sea animals: whale, shark, octopus or jellyfish) or inanimate (tools: screwdriver, saw, hammer or drill and instruments: flute, saxophone, violin or guitar) objects. All stimulus images were retrieved from the Google image search engine using the CCBY 4.0 filter. Half of the images were presented during the practice session (the first two of each subcategory mentioned), while the remaining were used for the experiment. Besides, each image was framed with a colored (green, pink, orange or blue) shape (triangle, circle, rhomboid or square).

The instructions gave information about the specific features of either one or the two target stimuli that had to be selected or integrated to respond. Task Demand specified whether the condition of the statement applied to both target stimuli (integration) or whether one of them had to be selected while ignoring the other (selection). Regarding the variable of Target Category, instructions pointed to the identity of the upcoming stimuli, with each regular trial containing either two animate or two inanimate objects. Each of the targets in a stimuli pair belonged to a different subcategory (animate trials: land animal and sea animal; inanimate trials: tool and musical instrument). Participants were not made aware of these constraints about the stimuli pairs. To ensure that participants were encoding all the information of the instructions (e.g., processing both target identities), we included 12,5% of *catch* trials (see e.g. González-García et al., 2021; Formica et al., 2021, for a similar approach) where a novel, non-instructed image was shown as target (4 trials per block, images could be of different target categories). Participants were told to detect those trials by pressing both keys simultaneously (“A” and “L”).

The variable of Target Relevant Feature indicated the specific features of the stimulus frames to pay attention to, either color (pink, green, blue, orange) or shape (square, rhomboid, circle, triangle). Finally, the instructions indicated the motor responses of either a left or right index button press. Half of the instruction sets referred to integration statements, whereas the remaining referred to selection statements; they were also equivalent in the rest of the parameters described (animate-inanimate stimuli; color-shape features), see Figure 1 for an example of the paradigm. Following this logic, to create the instruction set we combined 2 Task Demands (integration/selection) x 2 Target Categories (animate/inanimate) x 8 pairings of the animate or inanimate stimuli (each target of the pair belonging to a different subcategory) x 8 Relevant Features (4 colors/4 shapes) x and 2 response mappings (left or right button press if the stimuli fulfill the instruction), resulting in a total set of 512 novel instructions.

**Figure 1.**
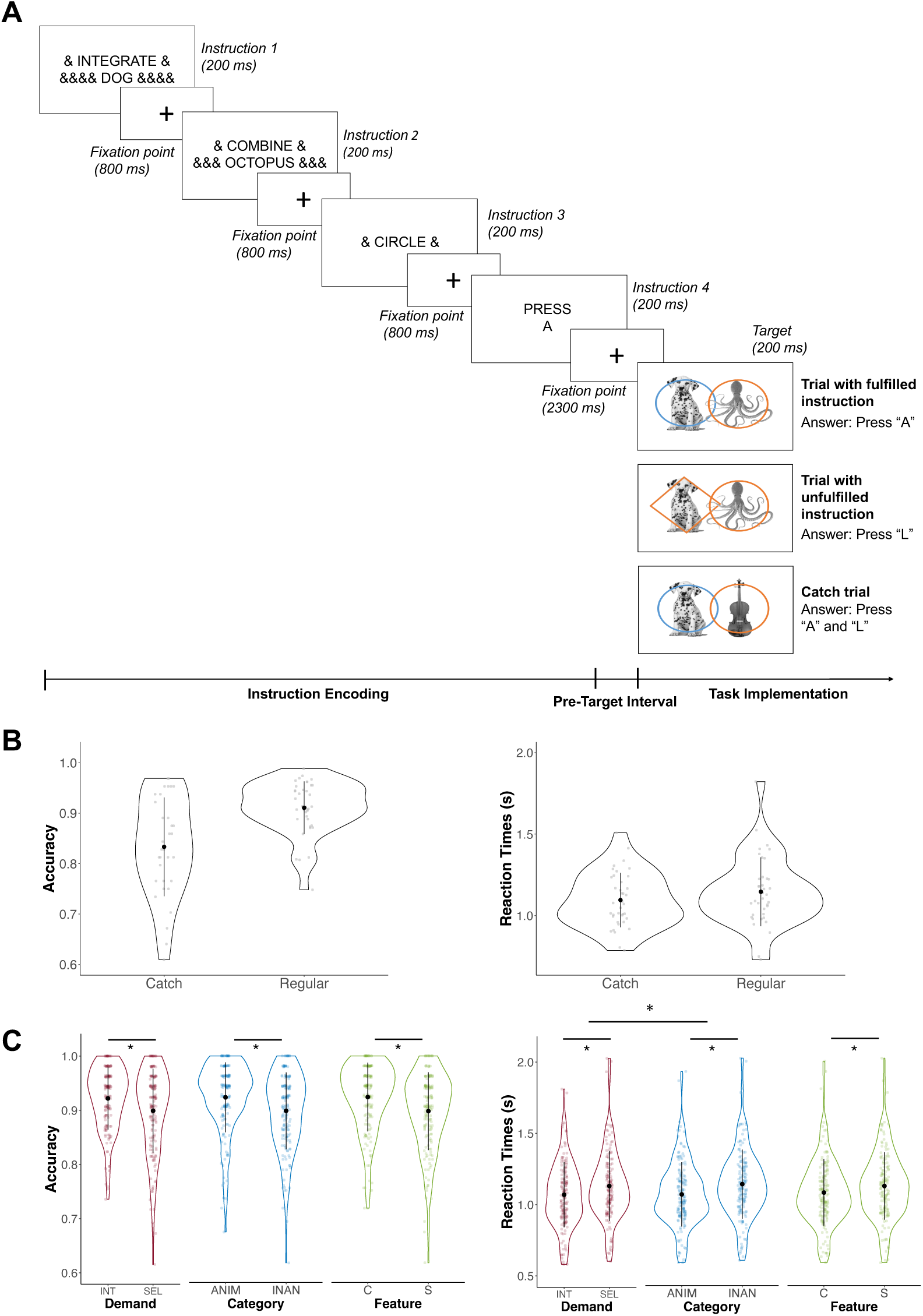
Task paradigm and behavioral results. **(A)** Example of a sequence of trial events. In this case, the sequential instructions indicate that if both the dog and octopus are framed by a circle, the key “A” has to be pressed. Trial examples for different cases and their correct responses are illustrated, instruction fulfilled (above), not fulfilled (middle), and catch trial (below). The original instructions were presented in Spanish. **(B)** Mean Accuracy (left) and Reaction Times (right) of catch and regular trials. **(C)** Repeated-measures ANOVA results for Accuracy (left) and Reaction Times (right) per condition. Black dots refer to the mean of the condition, and the black error bars span one standard deviation above and below the mean. Colored dots depict the mean per condition of every participant. Asterisks indicate significant main effects of the ANOVAs (p < 0.01). Abbreviations: INT, Integration; SEL, Selection; ANIM, Animate; INAN, Inanimate; C, Color; S, Shape.

To minimize horizontal eye movements while reading during EEG recording, the instructions were sequentially presented (see Figure 1A). The first and second screens indicated if the information from both stimuli had to be integrated (“Integra” and “Combina”, respectively, which translate to “Integrate” and “Combine”; ∼ 20° x 3.5°); or if the information of only one stimulus had to be selected (“Atiende” and “Ignora”, respectively, which translate to “Attend” and “Ignore”; ∼ 20° x 3.5°). Note that to match the presentation structure across task conditions, each Task Demand condition uses two different labels. They also included the two concrete stimuli that the instruction applied to (belonging to either animate or inanimate categories; ∼ 20° x 3.5°). The third screen indicated the specific feature to focus on, either a color (“Rosa”, “Verde”, “Naranja” or “Azul”; pink, green, orange, or blue, respectively; ∼ 20° x 3.5°), or a shape (“Cuadrado”, “Círculo”, “Triángulo” or “Rombo”; square, circle, triangle, or rhomboid, respectively; ∼ 20° x 3.5°). The fourth and last screen specified the motor response associated with the statement (“Pulsa A” or “Pulsa L”; which translates to “Press A” and “Press L”; ∼ 9° x 3.5°). For example, an instruction for the integration-animate-shape condition would be “Integrate dog, combine octopus, circle, press A”; here, the participant had to pay attention to both the dog and the octopus and if they were both framed by a circle, they had to press A (otherwise, L). An example of a selection-inanimate-color condition would be “Attend drill, ignore guitar, orange, press L” which would indicate “attend to the drill, ignore the guitar. If the drill is surrounded by an orange shape, press L, otherwise press A”. To ensure that on all trials each possible instruction had the same size on screen (first, second and third instructions, separately), meaningless symbols were added (‘&’) to match the number of characters (see Figure 1A).

The experiment presentation and behavioral data collection were done with Psychtoolbox running on Matlab on a Microsoft PC screen of 61 x 34 cm. Participants were seated at approximately 65 cm from the screen.

### 2.4 Procedure

Once participants arrived at the lab, they received overall instructions and performed a practice session. Only participants obtaining a minimum of 80% behavioral accuracy, for a maximum of 8 blocks, were invited to the EEG session (80% of participants reached this criterion). Practice sessions lasted from 10 to 40 minutes. Afterwards, we set up the EEG cap and participants performed the experiment that lasted approximately 1 hour and 40 minutes. Instruction-following and localizer blocks were interspersed throughout the session. The main task blocks were further differentiated according to the particular stimulus features that participants responded to (e.g., orange and blue frames in half of the blocks and pink and green frames in the other half). The whole experiment had a total of 24 blocks with self-pacing rest times between them. Of these, 16 blocks were of the main task and 8 of the localizer. To control for the effect of block order, the transitions between block types (main task with features 1, main task with features 2, and localizer task) were counterbalanced within participants, so that all transitions were equally probable. In total, the whole experimental session lasted approximately 3 hours.

Figure 1A displays a schematic representation of a trial of the main task. Each one of the four screens was on display for 200 ms and was followed by a fixation point for 800 ms. After the 4 screens (with their respective succeeding fixation intervals), participants had an additional 1500 ms pre-target interval. Then, the targets were shown for 200 ms (∼ 16° x 10.5°), with participants being instructed to respond as accurately and fast as possible. Participants had a 2800 ms interval afterwards to respond. This way, trials were divided into three phases: instruction encoding, pre-target and task implementation.

Each of the 16 blocks of the task contained 32 trials, giving a total of 512. For each block, there were 4 observations per each of the eight experimental conditions, resulting from the full crossing of the three independent variables. Additionally, for each block, we balanced the experimental conditions against a series of control variables that enabled creating a rich instructions pool, such as the target identity being integrated or selected (e.g., cat, drill…), the congruency of the target in regards to the instruction (both fulfilled it, both did not fulfill it, or only one fulfilled it), the specific frame features that participants responded to (e.g., green, square…), the response indicated in the instruction (e.g., press A, press L), the response required by the targets, and the font and format of the different instruction cues. Trial order was pseudo-randomized against these control variables, with sequential independence confirmed through a mutual information algorithm (e.g. González-García et al., 2021). This approach ensured no statistical dependencies among the experimental conditions included in the analysis and additional task manipulations of non-interest.

### 2.5 Behavioral Analyses

A 2×2×2 within-subjects repeated-measures ANOVA was performed, with the three independent variables of the task as factors: Task Demand (integration, selection), Target Category (animate, inanimate), and Target Relevant Feature (color, shape). This analysis was performed independently for behavioral Accuracy and Reaction Times using R on Rstudio (RDC Team, 2016). To investigate significant interaction terms, we conducted additional post hoc and applied a Holm correction to account for multiple comparisons.

### 2.6 Electrophysiological Analyses

#### EEG Data Acquisition and Preprocessing

Electrophysiological (EEG) activity was collected with BrainVision’s actiCAP equipment with 64 active electrodes. Of these, one was set as a reference channel (FCz) and two were used as electro-ocular channels (TP9, TP10). The electrodes’ impedance was maintained below 10 kΩ, as the amplifier’s manufacturers indicate. The EEG signal was recorded at a 1000 Hz sampling rate, and data files were structured following EEG-BIDS (Pernet et al., 2019).

EEG was preprocessed with the EEGLAB toolbox in Matlab (Delorme & Makeig, 2004) and custom Matlab scripts (López-García et al., 2022; available at https://github.com/Human-Neuroscience/eeg-preprocessing). EEG recordings were down-sampled at 256Hz, filtered with a low-pass at 126 Hz, a high-pass at 0.1 Hz, and an additional notch to remove the effect of the electrical current and its harmonics (49-51 Hz and 99-101 Hz). We generated epochs including the whole trial, covering a time window of 7200 ms (from -200 ms to 7000 ms). Epochs were locked at the onset of the first instruction and spanned until the end of the inter-trial interval after target presentation. After visual inspection, noisy channels were not included for the following steps (only one channel from one participant was removed). An Independent Component Analysis (ICA) was used to remove ocular (blink and saccades) and muscular artifacts identified through visual inspection and ICLabel (Pion-Tonachini et al., 2019). A mean of 2.87 components were removed per participant, ranging between 1 and 5. Then, an automatic rejection process was performed by removing trials with non-stereotypical artifacts that were not excluded previously with ICA. Reasons for exclusion were: 1) extreme values of voltage, in which the amplitude exceeded a range of ±150 µV; 2) abnormal spectra, in which the spectrum differed significantly from the baseline in the 0-2 Hz and 20-40 Hz frequency windows, linked to linear drifts in the signal caused by artifacts; 3) improbable data, with voltage values more than 6 standard deviations from the mean probability distribution. Noisy channels that contaminated the signal were removed and interpolated. The average reference was computed by pooling all the channels and then re-referencing to the mean. Finally, epoched data were baseline-corrected (-199 to 0 ms). Only correct and non-catch trials were included in the analyses, with an average of 44.60 ± 5.80 trials per condition (e.g., integration-animate-shape) and participant.

#### Representational Similarity Analysis (RSA)

The extent to which activity patterns were structured according to the instructed task components was evaluated through multivariate Representational Similarity Analysis (RSA; Kriegeskorte, 2008). By calculating the pairwise distance across all conditions, this analysis enables the abstraction of the neural data into a common representational space. A time-resolved RSA was performed to obtain a fine-grained distribution of the neural patterns throughout the entire trial. The analysis procedure was adapted from Peñalver et al. (2023) and was performed as follows:

To construct the empirical Representational Dissimilarity Matrices (RDMs), we crossed all the levels of the three independent variables (Task Demand: integration, selection; Target Category: animate, inanimate; Target Relevant Feature: color, shape). This yielded a total of 8 experimental conditions. For every condition and participant, EEG data was selected and averaged every 3 time points along the entire trial epoch (-200 to 7000 ms) to reduce the processing load. Voltage values were z-score normalized across trials. To calculate the distance between each pair of conditions, a cross-validated measure of Pearson’s correlation coefficient was employed. This cross-validated approach diminishes the risk of bias in the results (Grootswagers et al., 2018; Walther et al., 2016). For every participant, a 3-fold cross-validation went as follows: first, data of every participant was randomly divided into 3 chunks, matching condition sizes. On each of the 3 cross-validation folds, one chunk was assigned as the test set, while the remaining 2 as the training set. On every fold, the vectorized patterns of neural activity per condition were trial averaged and centered around 0 by subtracting the mean across conditions of each electrode. Per participant and time point, Pearson’s correlation was calculated for every pair of conditions between the training and test sets. The resulting matrices were transformed into distance matrices by computing 1 – Pearson’s coefficient (2 as more dissimilar and 0 as more similar). This was repeated until all chunks had been part of the test set, and the distance values were averaged across the 3 cross-validation folds. An empirical 8 x 8 Representational Dissimilarity Matrix (RDM) was thus obtained per subject and time point.

Theoretical RDMs were created to reflect the expected distances between our conditions according to the variables of interest (Figure 2A). We built three theoretical models driven by the relationship between the instructed components: (1) Task Demand, (2) Target Category, and (3) Target Relevant Feature. All matrices were fully orthogonal to each other. To estimate the share of variance that was explained by each theoretical model at every time point, we fitted them into a multiple linear regression. The theoretical RDMs were included as regressors and the empirical RDM from a given time point as the dependent variable. All matrices were vectorized before performing the regressions. To avoid illusory effects, the diagonal was removed from both empirical and theoretical RDMs (Ritchie et al., 2017). Since the cross-validated approach by its nature does not yield a fully symmetric matrix, all the remaining off-diagonal elements of the matrices were included in the regression. For each participant, we obtained a t-value of every model per time point, where every portion of the variance explained was unique to one specific model.

**Figure 2.**
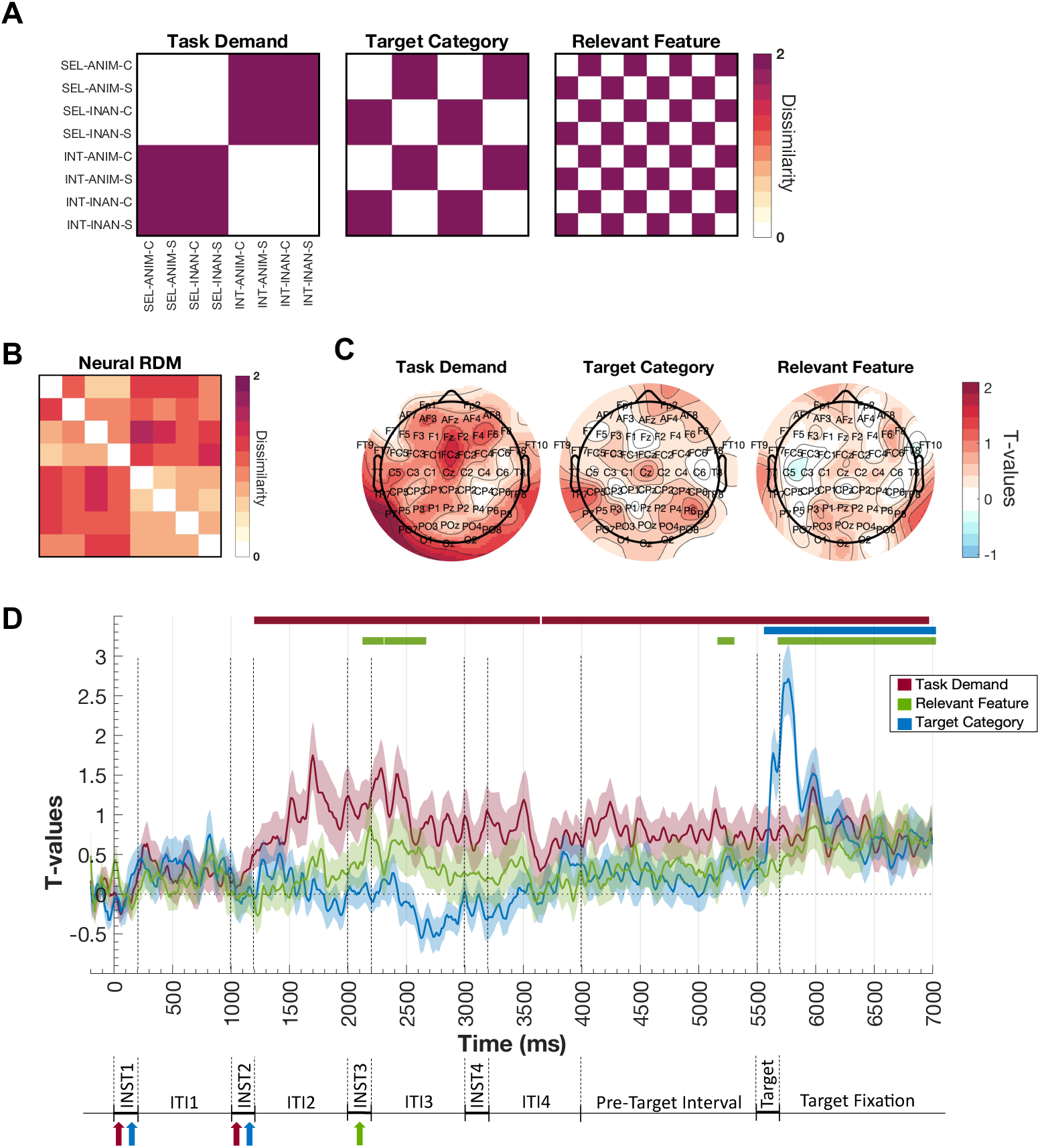
Task-relevant information organizes neural coding space: Representational Similarity Analysis (RSA). **(A)** 8×8 RDM of the theoretical models for the RSA of Task Demand, Target Category, and Target Relevant Feature. As the color bar illustrates, darker colors refer to maximal dissimilarity (2), and lighter colors to minimal dissimilarity (0). **(B)** Example of a neural RDM of one participant at one time-point. **(C)** RSA results per channel of the Task Demand, Target Category, and Target Relevant Feature model, representing the electrode’s topography of each model for visualization purposes. We performed Representational Similarity Analysis separately for each channel, using the time points as features. The empirical matrices per channel were fitted into a linear regression with the theoretical models as regressors. Colors refer to different t-values, with red illustrating higher t-values. **(D)** RSA time-resolved results. Each darker-colored line represents the t-values after fitting the three theoretical models with a multiple linear regression, illustrating the unique share of variance explained by each model. Lighter shading refers to the standard error associated with the mean t-values. The horizontal-colored lines above refer to the statistical significance against zero (dotted line) after implementing cluster-based permutation analysis. The results are accompanied by a simplified illustration of the task paradigm, where colored arrows indicate in which instruction’s screen the information of each task component is provided. Abbreviations: INT, Integration; SEL, Selection; ANIM, Animate; INAN, Inanimate; C, Color; S, Shape; A; INST, Instruction; ITI, Inter-instruction Interval.

Statistical significance was assessed non-parametrically via cluster-based permutation (Maris & Oostenveld, 2007; López-García et al., 2022). At the individual level, the labels of the theoretical RDMs were randomly permuted and the permuted matrices were included as regressors on a linear regression. This process was repeated 100 times per participant, resulting in 100 chance level t-values per theoretical model and participant. Next, we created 10^5^ group-level null t-value curves by randomly selecting individual subject permutation results and averaging across them. These permuted maps were used to estimate the above and below thresholds of the empirical chance distribution, which considered the 95-percentile of the distribution at each time point and the 95-percentile cluster size detected of contiguous time-points. To determine the smallest cluster size deemed significant for alpha = 0.05, multiple comparisons’ correction was implemented through the False Discovery Rate (FDR).

#### Time-resolved Multivariate Pattern Analysis (MVPA)

We used time-resolved MVPA (Grootswagers et al., 2018) to infer whether and when the EEG activity patterns encoded the three task-relevant components manipulated by our variables. We trained and tested the classifiers to distinguish between the two levels of each independent variable separately: integration vs. selection, animate vs. inanimate, and color vs. shape trials. Classifications were done on epochs (0 - 7000 ms) that included both task preparation (during instructions’ presentation and pre-target windows) and implementation (after target onset). Pre-processed raw voltage values for every channel and time point were used as features. All analyses were performed with the MVPAlab Toolbox (López-García et al., 2022; available at https://github.com/dlopezg/mvpalab). To minimize computational costs, the classifications were performed every 5 time points. Activity patterns were normalized across trials based on the standard deviation (King and Dehaene, 2014), computed in the cross-validation loop. Both training and testing sets were standardized within each fold in the following manner:

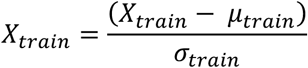

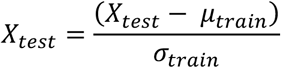

The *μ_train* and *σ_train* parameters reflect the mean and standard deviation of each feature (channels) of the training set. Next, we smoothed the data using a moving average filter with a sliding time window length of 5 time bins, to increase the signal-to-noise ratio (although slightly reducing the temporal resolution of the data). Afterwards, the number of trials across conditions was balanced to ensure an even distribution of the two classes.

A 5-fold cross-validation approach was followed. Briefly, the data were divided into five chunks and the algorithm used the first four as a training set while testing on the remaining one. This process was iterated five times, until all chunks were used as training and test sets. The final classifier’s performance value for each timepoint consisted of the mean performance across folds. Linear Discriminant Analysis (LDA) was used as a classification algorithm since it has shown good performance with EEG data while reducing computational costs (Grootswagers et al., 2018). The area under the curve (AUC) was the performance measure, as it is a non-parametric criterion-free estimate that has been recommended when dealing with 2-class classifications (King & Dehaene, 2014). AUC’s interpretation is similar to accuracy’s: in binary classifications, 0.5 equals chance level (equal probability of true and false positives) and 1 indicates perfect between-class segregation.

Statistical inference to detect above-chance classification was based on a non-parametric cluster-based permutation protocol (Maris & Oostenveld, 2007; López-García et al., 2022). The labels of trials were randomly permuted 100 times for each participant, obtaining 100 AUCs results that represented empirical chance level distributions. Afterwards, we created 10^5^ group-level null AUC curves by randomly selecting individual subject permutation results and averaging across them. Then, we employed these permuted maps to threshold our results considering both the 95-percentile null AUC value at each time point, and the 95-percentile cluster size of contiguous time points detected in the permuted null distribution. The final distribution was centered around an AUC value of 0.5 (chance level). Next, all the time points of the group’s permuted AUC maps that exceeded the estimated threshold were collected, yielding the normalized null distribution of cluster sizes. A False Discovery Rate (FDR) was implemented at a cluster level as a correction for multiple comparisons, allowing to achieve the smallest cluster size of contiguous time points deemed significant (López-García et al., 2022). This process was done independently for every variable analyzed.

#### Temporal Generalization Analysis

To examine whether the activity patterns coding the instruction components were stable or changed over time, and also to address if the classifiers generalized from preparation to task implementation, we performed a Temporal Generalization Analysis (King & Dehaene, 2014) for each of the independent variables. Following the parameters described above, we trained a classifier on a given time point and tested it on all the time points of the window. We iterated this protocol across the whole epoch, obtaining 3 temporal generalization matrices showing the AUC estimates for every train-test combination. To obtain statistical significance, the procedure indicated for time-resolved analysis was repeated (see *Time-resolved MVPA* in Methods), with the exception that individual-level permutations were reduced to 10 and group-level permutations to 10^4^, to decrease computational load.

#### Cross-Condition Classification Performance (CCGP)

The CCGP allows quantifying the extent to which the neural codes of the three instruction components support generalization, by examining through cross-classification all possible ways of choosing training and testing data (Bernardi et al., 2020). A high average CCGP suggests that the underlying neural codes support task-specific low-dimensional geometries of the neural space. In this case, the classifiers are generalizing across different train-test combinations of the conditions, and not only over noise related to trial-to-trial variations (Bernardi et al., 2020), but they do so at the cost of separability (Badre et al., 2021).

First, the train and test data were split according to the condition’s label, with the training and testing sets always consisting of different conditions. Considering that we had 3 main instruction components, the full-crossing of the variables resulted in 8 conditions. For the CCGP analysis of every instruction component (Task Demand, Target Category, and Target Relevant Feature), there were 4 conditions of one level of the variable and 4 of the other, creating a balanced dichotomy of the data (4 conditions of class A vs. 4 conditions of class B). We performed a cross-classification loop by setting 2 of the 8 conditions as the testing set (1 condition of class A vs. 1 condition of class B), while the remaining conditions were kept as the training set (3 conditions of class A vs. 3 conditions of class B), since larger training sets lead to better generalization performance (Bernardi et al., 2020). For example, to compute the CCGP of Task Demand, in each iteration of the analysis we trained a classifier to discriminate the neural patterns of integration and selection from 3/4 of the conditions (e.g., [Selection-Animate-Color], [Selection-Animate-Shape], [Selection-Inanimate-Shape] vs. [Integration-Animate-Shape], [Integration-Inanimate-Color], [Integration-Inanimate-Shape]), and then tested it against the remaining 1/4 of the conditions (in this example, [Selection-Inanimate-Color] vs. [Integration-Animate-Color]). Then, this process was iterated for all unique combinations of the conditions into training and testing sets, yielding a total of 16 repetitions (Bernardi et al., 2020, see Figure 4–1 in Extended Data). Hence, classifiers were always tested on neural patterns of different conditions from the ones they were trained on. Given that the analysis was repeated along the 16 possible combinations, a 16-step cross-validation loop was implemented along the conditions. For this reason, CCGP analyses usually result in lower, although informative, performance scores than regular MVPA (Bernadi et al., 2020).

For every task variable separately, and every iteration of the possible training and testing sets, we performed time-resolved MVPA with a cross-classification approach (López-García et al., 2022). We implemented the same protocol as in *Time-resolved MVPA*, the only change was that no additional cross-validation of the trials was implemented; since training and testing sets always came from different data, there was no risk of overfitting. The mean cross-classification performance after iterating through all 16 training and testing combinations of each task component was the Cross-Classification Generalization Performance (CCGP) of that variable.

To assess statistical significance, a non-parametric cluster-based permutation protocol was implemented (Maris & Oostenveld, 2007; López-García et al., 2022). Per subject and task variable, the labels of the conditions of each train-test combination were randomly permuted 10 times, resulting in 10 AUCs maps, which represented the empirical chance level distribution of each train-test combination. Then, the permuted maps were averaged across all 16 possible train-test combinations, obtaining 10 permuted maps per subject and instruction component. To perform group-level statistics and obtain the significance of the results, the same steps were followed as in *Time-resolved MVPA*. Significant above-chance AUCs reflect that the decoded task components were coded in an abstract format that favors generalization.

#### Correlation between Time-Resolved MVPA and CCGP

To test whether the temporal patterns of regular Time-resolved MVPA and CCGP were similar, suggesting that when the task components were encoded in the neural patterns there was matching evidence of abstract coding of information, we performed Pearson’s correlations. Per participant, we calculated the Pearson’s correlation coefficient (*r*) of the MVPA and CCGP results including all time-points of the trial, separately for each task component. To compute the average coefficient across participants, we used Fisher’s z-transformation (formula below; Fisher, 1970) to create a normal distribution of the correlation values before averaging and then reverted to Pearson’s correlation coefficient (second formula below).

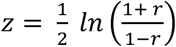

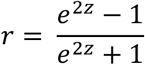

To compute the average p-value of the correlations across participants, we used Fisher’s method (formula below; Fisher, 1970). By summing the log-transformed p-values, we obtained Fisher’s statistic (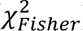), which follows a chi-square distribution. To revert to p-values, we calculated the Complementary Cumulative Distribution Function of Fisher’s statistic, which provides the upper tail probability of the chi-square distribution.

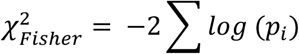

To ensure that the correlation coefficients obtained were robust and not due to the temporal structure of the noise, we repeated this analysis comparing the regular MVPA and CCGP results of different variables pair-wise (e.g. correlation between MVPA results of Task Demand and CCGP results of Target Category), yielding 6 control comparisons.

#### Temporal Generalization Analysis with cross-classification

To extend the information given by the CCGP, we used Temporal Generalization Analysis with a cross-classification approach. We thus addressed whether the generalizable neural patterns of the instruction components were also stable through the entire epoch. This was performed separately for every instruction component and across the different levels of the remaining variables. In other words, the classifier had to discriminate between the two levels of a variable when trained and tested on different contexts, to establish if the neural codes were generalizable with independence of the condition of the other variables. To reduce computational load, we did not explore all possible data splits of the variables, but pairwise. This way, the analysis evaluated whether Task Demand was generalizable across different Target Categories, and across different Target Relevant Features; whether Target Category was generalizable across different Task Demands, and across different Target Relevant Features; and whether Target Relevant Feature was generalizable across different Task Demands, and across different Target Categories. The analysis protocol was the same as in *Temporal Generalization Analysis*, with the only difference being that training and testing data always came from different conditions. The analysis was done in both directions of the data, i.e. when trained on condition A and tested on condition B, and vice-versa. Afterwards, the mean results and permutation maps across both directions were obtained, and statistical group analysis was performed for each direction and the mean separately.

#### Dimensionality Analysis

The extent to which the neural activity generated by the novel instructions is coded in a high-dimensional space was tested through a decoding-based dimensionality analysis that measured the decodability of all the binary dichotomies of the conditions with a linear classifier (Bernardi et al., 2020). The number of decodable dichotomies after performing all possible mixtures of the instruction conditions provides information about how many “dimensions” can be extracted from the neural data (Rigotti et al., 2013; Fusi et al., 2016). A high number of decoded dichotomies implies more separable or expanded codes, reflecting higher dimensionality, while a low number implies more compressed neural codes in a lower dimensional space (Rigotti et al., 2013; Fusi et al. 2016).

We followed an analysis protocol similar to Bernardi et al (2020). Given that crossing of our three instruction variables yields a total of 8 conditions, we obtained 35 possible dichotomies with a balanced number of conditions per side of the classification (4 vs. 4; see Figure 5-1 in Extended Data). Out of the 35 dichotomies, three correspond to our main instruction components, while the rest depict a mixture of the variables. Thus, each mixed dichotomy represents a complex combination of the three task variables, which hinders interpretation of results per dichotomy. For every dichotomy of the data, we performed a MVPA analysis as indicated in *Time-resolved MVPA*. Hence, the linear classifier was trained and tested to discriminate between the two sides of the dichotomy. This analysis was repeated per time-point of the trial epoch.

We calculated two complementary dimensionality measures: the number of dichotomies decoded above-chance and the Shattering Dimensionality index (SD; Bernardi et al., 2020). First, to calculate the number of decoded dichotomies (Rigotti et al., 2013; Bhandari et al., 2024), we assessed how many dichotomies were significantly decoded above chance per time point. To obtain this, we performed a non-parametric cluster-based permutation protocol (previously explained; Maris & Oostenveld, 2007; López-García et al., 2022) at the level of each dichotomy. This way, we obtained the instances of the epoch that had been significantly decoded per dichotomy. Afterwards, we calculated the total number of significantly decoded dichotomies by adding the dichotomies of above-chance classification per time-point.

We also calculated the Shattering Dimensionality index (Bernardi et al., 2020) as the mean decoding performance across all possible dichotomies. Although this averaged index could over-represent dichotomies with higher classification accuracy, we included this measure following past studies (Bernardi et al., 2020; Bhandari et al., 2024; Posani et al., 2024) to further qualify the above-mentioned dimensionality count estimates. To infer statistical significance, we again implemented a non-parametric cluster-based permutation protocol (Maris & Oostenveld, 2007; López-García et al., 2022). The labels of the conditions were randomly permuted 10 times per participant and dichotomy, to obtain the empirical chance distribution of each dichotomy. Afterwards, the permuted maps were averaged across all 35 dichotomies, yielding the permuted maps per subject. After group-level statistics were implemented on the averaged decoding performance (see *Time-resolved MVPA* in Methods), significant above-chance AUCs reflected the Shattering Dimensionality index, with separable neural codes of the data.

#### Neural Geometry - Representational Similarity Analysis (RSA)

We further tested the geometrical arrangement of patterns of brain activation across the two task demands instructed, integration vs. selection of information, to study whether their coding structure favored generalization across other instruction constituents while keeping the two task contexts disaggregated. To estimate the neural geometries between trials with different task components, we employed Representational Similarity Analysis (RSA; Kriegeskorte, 2008, following the approach of Muhle-Karbe et al., 2023). We measured the proportion of variance in the neural empirical matrices explained by two geometrical models: offset and orthogonal (see Figure 6A). The theoretical models represented two hypotheses about how Task Demand, our highest-level variable, was geometrically encoded over a lower-dimensional manifold. The empirical RDMs were calculated following the same steps as in *RSA*; thus, 8×8 neural matrices per subject and time point were obtained.

The theoretical RDMs were built by estimating the pairwise distances between the conditions according to two geometry hypotheses about the spatial configuration of the Task Demand: a) the Offset Model is in line with the two levels of Task Demand (integration and selection) being represented in low-dimensional space as separable and parallel planes, which favors generalization but is prone to interference; b) the Orthogonal Model aligns with coding where integration and selection are represented in a geometry that impairs generalization but prevents cross-task interference, with the configuration between both conditions creating a 90° angle (see Figure 6A). The values determined for the generated geometry RDMs ranged from 0 (similar) to 2 (dissimilar) and the matrices included the full crossing of the 3 task components (Task Demand, Target Category, and Target Relevant Feature), resulting in 8×8 RDMs.

To investigate the variance explained by each geometry model, they were fitted into a multiple linear regression. The geometry models were included as regressors, while the neural RDMs were the dependent variables, with all matrices vectorized before conducting the regression. Thus, we obtained the t-values unique to each model. This was repeated separately per participant and time point. Finally, statistical significance was obtained via non-parametric cluster-based permutation testing (see *RSA* in Methods; Maris & Oostenveld, 2007; López-García et al., 2022).

## 3. Results

### Behavioral results

Participants were able to perform the task with efficiency, as shown in Figure 1B. Regular trials had a mean Accuracy of 0.91 (SD = 0.05) and mean Reaction Times of 1.13 seconds (SD = 0.22). Catch trials also add a good overall performance, with a mean Accuracy of 0.780 (SD = 0.16; Reaction Times: M = 1.09 s, SD = 0.18), which confirms that participants were attending to all the information given by the instructions.

We explored whether the three main task components modulated behavior. The two behavioral measures (Accuracy and Reaction Times) in regular trials were analyzed separately with two within-subjects repeated-measures ANOVAs, including as factors Task Demand (integrate or select information), Target Category (animate or inanimate), and Target Relevant Feature (attend to the color or shape surrounding the images). The results can be seen in Figure 1C. Task Demand had a significant main effect on both Accuracy (*F*_38,1_ = 9.87; p < 0.01; 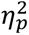 = 0.21) and RT *(F*_38,1_ = 30.51; p < 0.01; 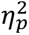 = 0.45), with more accurate (integration: M = 0.92, SD = 0.06; selection: M = 0.90, SD = 0.08) and faster responses (integration: M = 1.07, SD = 0.23; selection: M = 1.13, SD = 0.24) for integration trials. Target Category also showed a significant main effect on Accuracy (*F*_38,1_ = 25.91; p < 0.01; 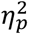 = 0.41) and RT (*F*_38,1_= 86.67; p < 0.01; 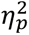 = 0.70), with more accurate (animate: M = 0.92, SD = 0.07; inanimate: M = 0.90, SD = 0.07) and faster responses (animate: M = 1.06, SD = 0.23; inanimate: M = 1.14, SD = 0.24) for animate trials. Lastly, we also found a significant main effect of Target Relevant Feature on Accuracy (*F*_38,1_ = 27.52; p < 0.01; 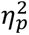 = 0.42) and RT (*F*_38,1_ = 70.32; p < 0.01; 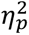 = 0.65), with more accurate (color: M = 0.92, SD = 0.06; shape: M = 0.90, SD = 0.07) and faster responses (color: M = 1.08, SD = 0.24; shape: M = 1.12, SD = 0.24) for color trials. Additionally, we found a significant interaction term between Task Demand and Category on Reaction Times (*F*_38,1_ = 14.05; p < 0.01; 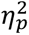 = 0.27). Post hoc tests showed that the effect of Task Demand was significant for both Category conditions, with a larger effect for inanimate (*t*_77_ = -7.66; p < 0.01) than animate trials (*t*_77_ = -4.42; p < 0.01). All the remaining interactions in terms were not significant (p > 0.05). Thus, our findings reveal that the task components mostly had independent effects on behavior.

### Novel task information organizes the underlying neural space at different temporal dynamics

Multivariate model-based Representational Similarity Analysis (RSA; Kriegeskorte, 2008) allowed to compare the neural data with the theoretical RDMs of the three instructed variables and examine how well each model reflects the neural coding space along the entire trial epoch. Neural Representational Dissimilarity Matrices (RDMs; Figure 2B) were created from a cross-validated Pearson’s coefficient measure (1 - r), and later fitted into a multiple linear regression as dependent variables, with the theoretical RDMs of each of the three task components as regressors (Figure 2A). The time-resolved results showed that Task Demand, the higher-level task component, had a significant effect during most of the instruction stage while the remaining lower-level variables did so transiently and only in specific time windows, as displayed in Figure 2D.

### Novel task information is encoded in the neural patterns with fluctuating temporal stability during preparation and execution

To study the coding of content of the task components in neural patterns, we used binary classifiers with a Linear Discriminant Analysis algorithm independently for every variable (Task Demand, Target Category, and Target Relevant Feature). In line with the results of the Representational Similarity Analysis (RSA), the Task Demand could be decoded during most of the epoch (Fig. 3A). First, two early peaks were significant during the processing of the first instruction (starting at 760 and 950 ms; colored arrows at the bottom of Fig. 3A). Next, a wide significant cluster extended during the remaining of instruction presentation (for a 3000 ms window), most of the pre-target interval, and target processing. The highest Area Under the curve (AUC) values for Task Demand were found during the processing of the second and third sections. Second, the Target Category classifier performance was significantly above chance in two shorter clusters during the instruction phase, right after the presentation of the second and third instruction screens (for 500 and 150 ms windows, respectively), which were those indicating the relevant category and feature. There was a small ramping-up of the classifier’s precision right before the target onset (starting at 5250 ms) and the curve was steadily significant after target presentation (5540 ms onwards; Fig. 3A). Last, the classifier decoding of the Target Relevant Feature was significant right after the third instruction screen (for a 350 ms window; Fig. 3A), where the feature information was displayed. No significant decoding was detected during the pre-target interval or after target presentation.

**Figure 3.**
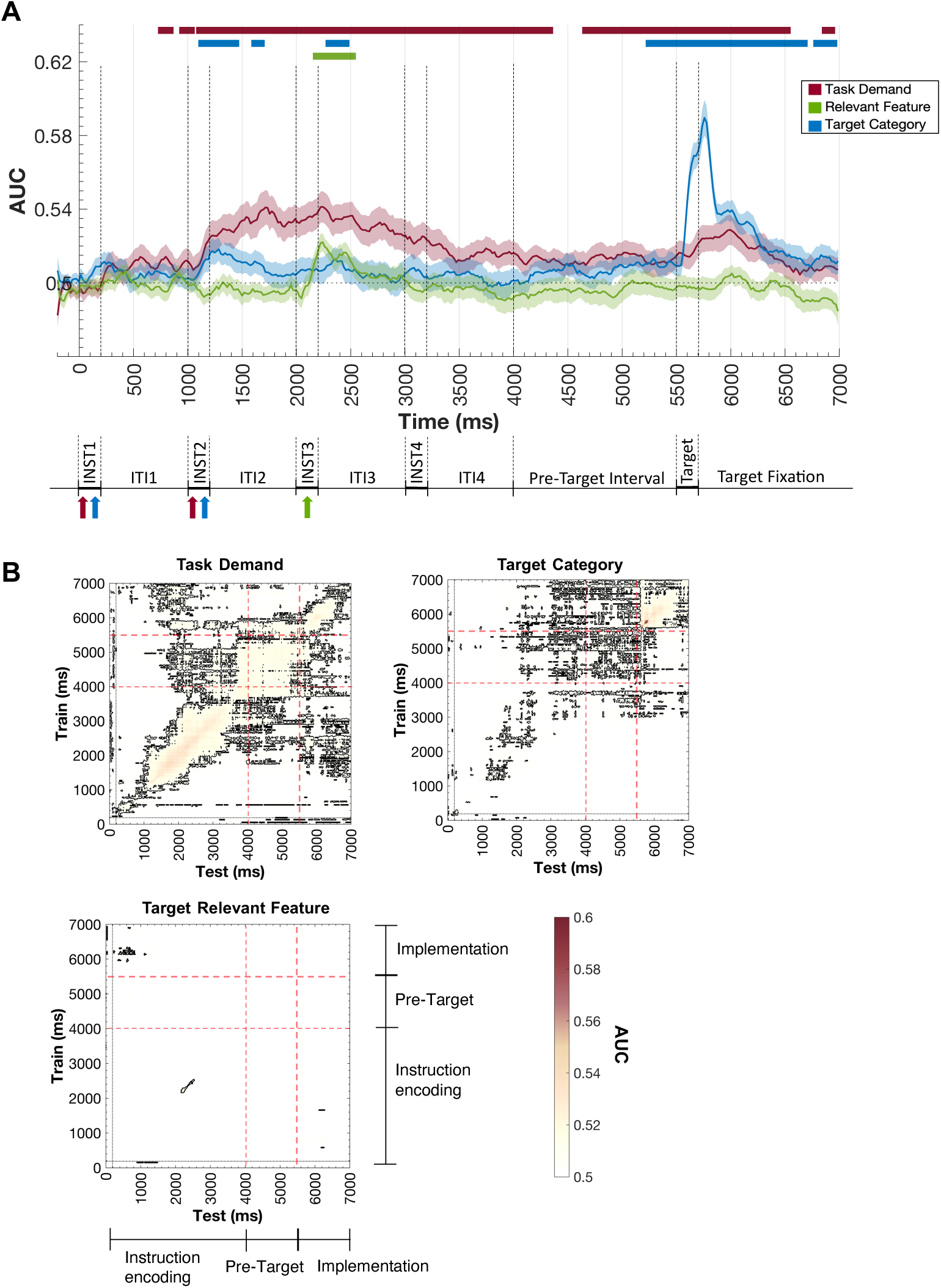
Raw Voltage MVPA Results **(A)** Results of the time-resolved classifications of the instruction-following task. Darker thin lines illustrate Area Under the Curve values (AUC), and lighter shading refers to the standard error associated with the mean (SEM). The horizontal-colored lines refer to the statistical significance against chance (dotted line). The figure shows the results of training and testing as described in *Time-resolved MVPA* (Methods section). The results are accompanied by a simplified illustration of the task paradigm. **(B)** Results for the temporal generalization analyses: Task Demand, Target Category, and Target Relevant Feature Temporal Generalization Matrices. The vertical axis indicates the training time bins, and the horizontal axis the testing time bins. The color bar indicates the AUC value of the classifier, and black lines denote significant clusters. Red dotted lines indicate the partitions between instruction encoding, pre-target interval, and task implementation. Abbreviations: INST, Instruction; ITI, Inter-instruction Interval.

The preceding findings do not address the degree to which neural coding patterns are stable over time. To further examine whether common or independent task representations are recruited across the preparatory and implementation stages, we used a temporal generalization approach (King & Dehaene, 2014). The classifier was trained at one time point and tested on all the time points of the epoch, allowing the comparison of different patterns of brain activity throughout the entire trial (see Figure 3B).

First, regarding Task Demand (Fig. 3B upper-left) we observed off-diagonal significant classification, reflecting the sustained activity of the same patterns across time. Two connected clusters of significant generalization during the encoding of instructions (1000-2000 and 2000-3000 ms) suggest common representations within these two intervals, with activity patterns during the pre-target interval also displaying generalization (4000-5500 ms; middle quadrant of the matrix). During the task implementation phase, there was a small cluster of significant generalization after the target presentation (6000-7000 ms; upper-right quadrant of the matrix). The cross-classification across different task stages shows small significant clusters with generalized activation between the encoding of the last components of the instructions (2000-4000 ms) and both pre-target (middle-left and bottom-medium quadrants of the matrix) and task implementation (upper-left and bottom-right) intervals, similar cross-decoding was also found across the pre-target and implementation stages (upper-medium and middle-right). However, the temporal profile of these findings does not follow a sustained pattern reflecting a complete temporal persistence of neural codes during the whole task trial. Rather, this suggests a certain degree of stability of part of the neural patterns coding across the three task stages.

As for the analysis of Target Category (Fig. 3B upper-right), transient epochs of significant decoding were detected during the presentation and processing of the instructions; without generalization across the different instructions’ screens. Several points of significant cross-classification were evident during the pre-target interval, but with no apparent unified cluster. This emerged after target onset (5500-7000 ms), indicating stable neural patterns during the task implementation period. Additionally, there were small clusters of significant cross-decoding when the classifier was trained during target processing (5500-7000 ms) and tested throughout instructions and pre-target intervals (upper-left and middle quadrants of the matrix). Besides, several significant points of cross-classification between the later instruction encoding and the pre-target interval indicated generalization between these two time periods. Overall, cross-classification results suggest that the animacy of the target category was coded with representations sustained to some extent. Last, the Target Relevant Feature matrix (Fig. 3B lower-left) only showed points of significant decoding on the diagonal during the encoding of the third instruction (2000-3000 ms).

### Abstract neural codes of different task-relevant information emerge with sustained and transient dynamics

After ascertaining that neural patterns code task-relevant information conveyed by the novel instructions, we addressed whether these patterns were represented in an abstract format, generalizing across different contexts. For that purpose, we performed Cross-Condition Generalization Performance (CCGP; Bernardi et al., 2020) for the three task variables in a time-resolved manner. We split the data into train and test sets for every possible combination of the conditions and performed linear classifiers, leaving 6 conditions for training (3 vs. 3) and 2 for testing (1 vs. 1) on every iteration, so that training and testing sets always consisted of different conditions. Figure 4B illustrates the CCGP value, the average performance of the 16 cross-classifications (see Figures 4-2, 4-3 and 4-4 in Extended Data for the classifier performance per cross-classification). Note that although the performance measure of the classifier had a generally low value compared with previous time-resolved MVPA analysis, this is an expected result when considering the cross-classification approach implemented (Bernardi et al., 2020).

**Figure 4.**
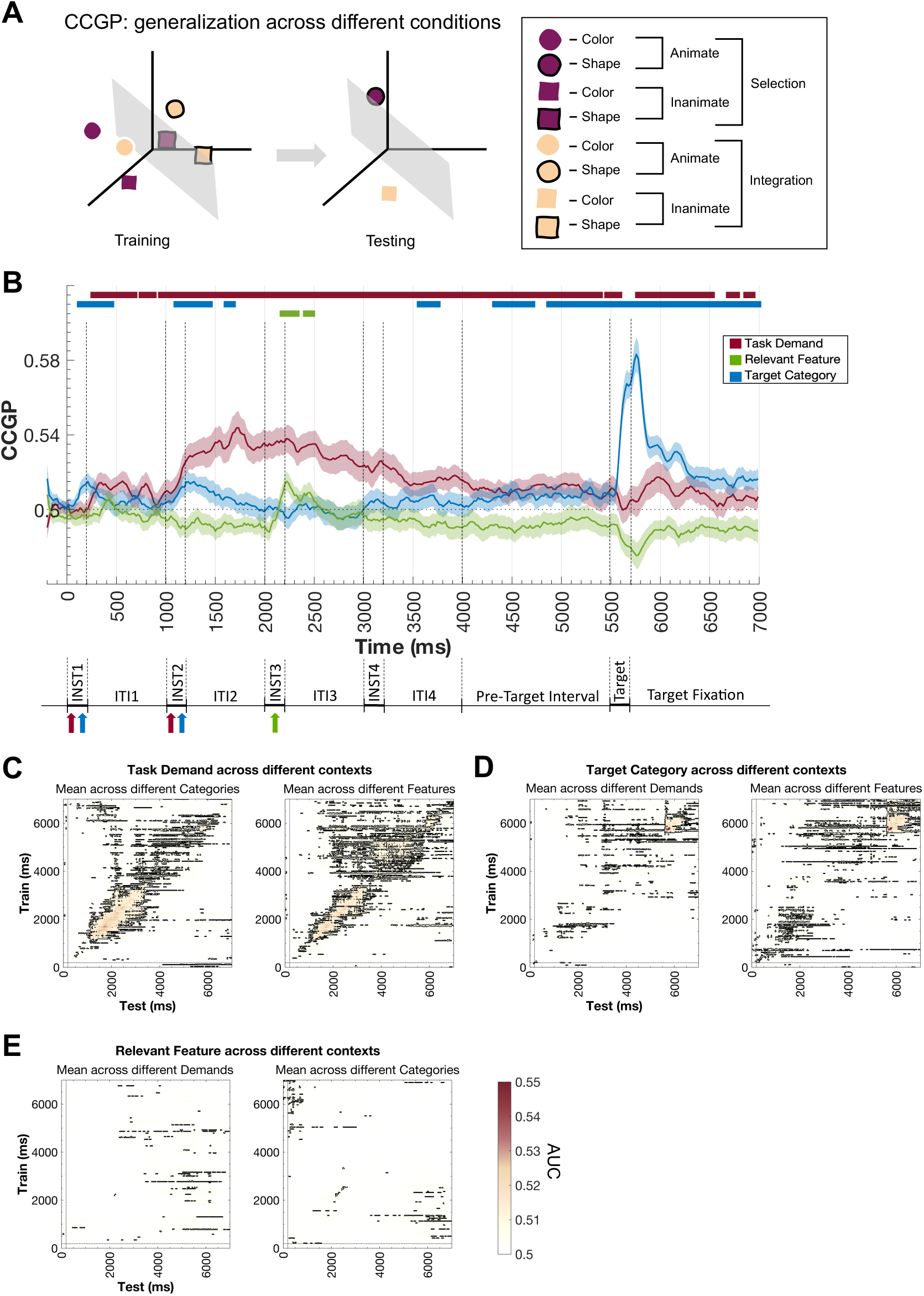
Temporal dynamics of abstract neural codes. **(A)** CCGP analysis rationale. Data were split according to the condition labels, so that training and testing sets always consisted of different conditions. This process was iterated for all possible splits of the data (see Figure 4-1 in Extended Data for full depiction). **(B)** Cross-Classification Generalization Performance (CCGP) of the task components. The darker-colored lines represent the mean Area Under the Curve (AUC) after performing a linear classifier over all possible combinations of train and test data sets for each variable: Task Demand (magenta), Target Category (blue), Target Relevant Feature (green). Lighter shading refers to the standard error associated with the averaged performance measure (SEM). The horizontal-colored lines refer to the statistical significance against chance (dotted line) after implementing cluster-based permutation analysis. The vertical axis indicates the training time bins, and the horizontal axis the testing time bins. The color bar indicates the AUC value of the classifier, and black lines denote significant clusters. **(C, D, E)** Results of Temporal Generalization analysis of the task components through different contexts with cross-classification (different conditions for train and test). **(C)** Mean Temporal Generalization Matrices after classifying integration vs. selection (Task Demand) through different Target Categories (left) and different Relevant Features (right). **(D)** Mean Temporal Generalization Matrices after classifying animate vs. inanimate (Target Category) through different Task Demands (left) and different Relevant Features (right). **(E)** Mean Temporal Generalization Matrix after classifying color vs. shape (Relevant Feature) through different Task Demands (left) and different Target Categories (right). See Figures 4-2, 4-3, and 4-4 in Extended Data for the MVPA results per dichotomy of Figure 4B. See Figure 4-5 in Extended Data for the correlation distributions across the pair-wise comparisons of regular MVPA and CCGP results. See Figures 4-6, 4-7, and 4-8 in Extended Data for the results per direction of the classification of Figure 4C, 4D, and 4E.

CCGP findings illustrate different temporal profiles of abstraction for each of the task components. The variable Task Demand engaged common neural codes along most of the trial, since the mean Area Under the Curve (AUC) of the CCGP was significantly above chance for almost all the time-points of the epoch. There was a peak of cross-generalized decoding during instruction encoding and, although the AUC decreased on the pre-target interval, it was still significantly decoded above chance. Thus, Task Demand recruited in a stable fashion common neural codes that support generalization, corroborating that the higher-level variable of the task was represented abstractly in a prolonged fashion. In fact, this temporal profile of results seems to concur with those of the regular Time-Resolved MVPA, suggesting that abstracted neural patterns underpin the coding of this task component throughout the trial window. Note that the cross-validation and cross-decoding schemes followed in these two analyses do not allow for direct statistical comparison due to the different number of observations used for testing and training the classifiers, which could lead to spurious results. Instead, we aimed at characterizing their potentially common temporal profiles through Pearson correlations, by including the results of all time points at once. The average coefficient across participants after Fisher’s Z-transform indicated a significant and strong relationship between the two time courses (*r_avg_* = 0.84, combined p-value using Fisher’s method, *p* < 0.01).

Target Category showed cross-generalization transiently during the earlier stages of the trial and steadier after target presentation, with a high CCGP peak right after target onset. The results suggest that Target Category was represented abstractly only at specific time intervals, which match the time periods where the content of this variable was decodable in the data. In this case, compared to the regular MVPA results, the average Pearson’s correlation coefficient across participants after Fisher’s Z-transform was *r_avg_* = 0.83 (combined p-value using Fisher’s method, p < 0.01). This suggests that whenever the Target Category was explicitly coded in the neural patterns, it was, at least partially, in a generalizable format. Lastly, the Target Relevant Feature showed a significant CCGP at a short interval during the encoding of the third instruction, when feature information was being provided, again matching the decoding results during this time period (see previous time-resolved MVPA results, average Pearson coefficient across participants was *r_avg_* = 0.77, p < 0.01). Thus, this portrays abstract transient coding of feature information.

To ensure that the correlation results were robust and not due to the temporal structure of noise, we repeated the correlation of regular decoding and abstraction results but with different variables (e.g., MVPA results of Task Demand and CCGP results of Target Category), for a total of 6 pair-wise comparisons. The results showed negative, close to zero correlation coefficients, with the average Pearson coefficients ranging from *r_avg_* = -0.10 to *r_avg_* = -0.17 across the 6 comparisons (p-values range from p = 0.11 to 0 = 0.19; see Figure 4-5 in Extended Data for the distribution of correlations across all pair-wise comparisons). Thus, the strong correlations from MVPA and CCGP of the same variable (described in the previous paragraphs) were not merely a by-product of the temporal noise structure in the data.

**Figure 5.**
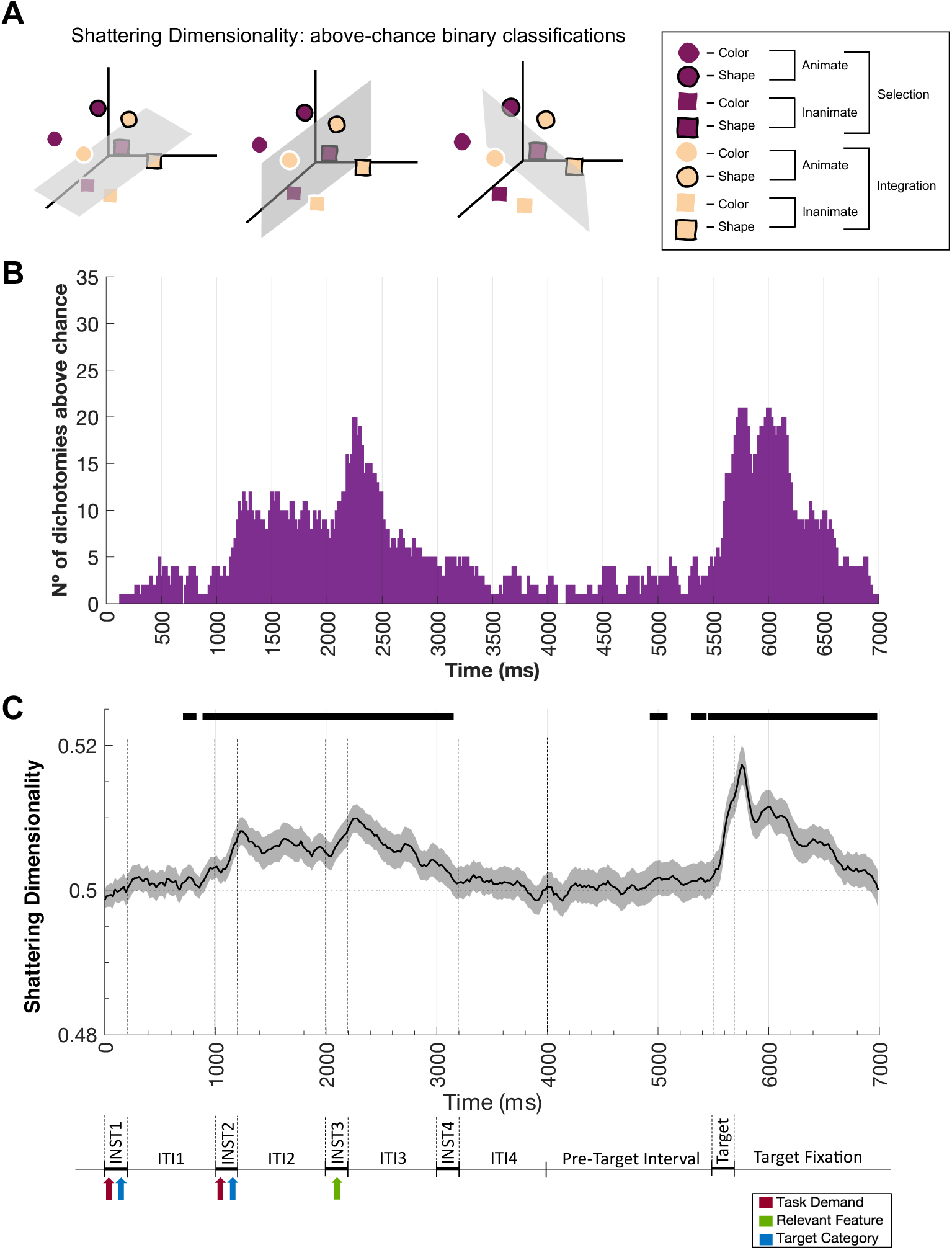
Temporal dynamics of neural dimensionality. **(A)** Analysis rationale. Data from the full-crossing of the variables were split into all possible balanced dichotomies (see Figure 5-1 in Extended Data for the full depiction of data splitting). After performing classifiers on every dichotomy, the more ways the data can be separated, the higher the dimensionality of the neural space. **(B)** Number of dichotomies decoded significantly above chance. A linear classifier was performed with each of the 35 balanced dichotomies obtained from mixing the task variables. For every time point, the significantly decoded dichotomies were summed. **(C)** Shattering Dimensionality (SD) results. The darker line represents the mean Area Under the Curve (AUC) after applying a linear classifier over all possible balanced dichotomies, obtained from mixing the task conditions. Lighter shading refers to the standard error associated with the mean of the performance measure. The horizontal line refers to the statistical significance against chance (dotted line) after implementing cluster-based permutation analysis. The results are accompanied by a simplified illustration of the task paradigm. See Figure 5-2 in Extended Data for the MVPA results per dichotomy. Abbreviations: INST, Instruction; ITI, Inter-instruction Interval.

To investigate whether these abstract neural codes were also temporally stable and transferable across the trial epoch, we performed a Temporal Generalization analysis with cross-classification. We obtained generalization matrices for each task variable when training and testing on different conditions of the remaining variables. The analyses were performed in both directions of classification (train on condition A and test on B, and vice-versa) and later averaged (see Figures 4-6, 4-7 and 4-8 in Extended Data for the results per direction of classification).

**Figure 6.**
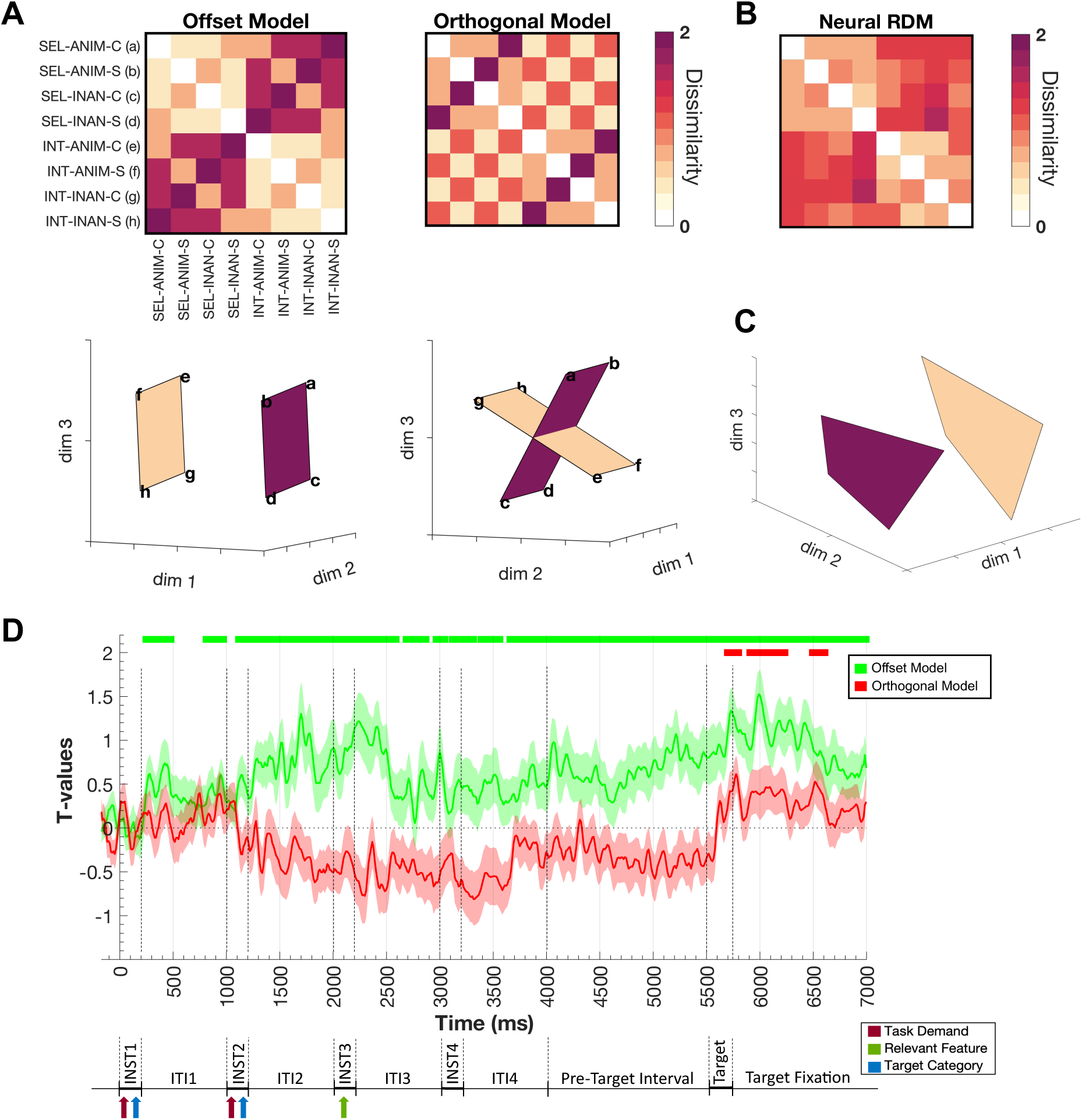
Representational Similarity Analysis (RSA) of neural coding geometry. **(A)** Representational Dissimilarity Matrices of the two geometry models and their Multidimensional Scaling plots (Offset and Orthogonal Model, respectively). The upper figures represent the theoretical RDMs; as the color bar illustrates, darker colors refer to maximal dissimilarity (2), and lighter colors to minimal dissimilarity (0). The figures below depict the two alternative geometries projected on a three-dimensional space via Multidimensional Scaling. The darker purple plane refers to the selection activity patterns (with each vertex indicating a condition, e.g., a = selection - animate - color), and the beige plane to the integration activity patterns. **(B)** Example of a Neural RDM of one participant at a specific time point. **(C)** Multidimensional Scaling (MDS) plot of the previous neural RDM reduced to 3 dimensions. MDS was used for visualization purposes only. **(D)** RSA results per geometry model. Each darker-colored line represents the t-values after fitting the two geometry dissimilarity models to a multiple linear regression, illustrating the unique share of variance explained by each model. Lighter shading refers to the standard error associated with the mean t-values. The horizontal-colored lines refer to the statistical significance against zero (dotted line) after implementing cluster-based permutation analysis. The results are accompanied by a simplified illustration of the task paradigm. Abbreviations: INT, Integration; SEL, Selection; ANIM, Animate; INAN, Inanimate; C, Color; S, Shape.

For the Task Demand we observed similar generalization patterns both across different Target Categories and Relevant Features (Figure 4C), with a unified cluster of generalization during the processing of the second and third instructions’ screens, and points of cross-classification prior to and after target onset. The only difference of note was a thinner instruction encoding cluster (fewer instances of cross-classification) across different Relevant Features (Fig. 4C right). In comparison with the regular temporal generalization results (with the same data for training and testing; see Temporal Generalization Analysis results), a smaller cluster of generalizable neural patterns during target processing was detected, which suggests that the Task Demand after the target appeared was coded with more independent neural patterns depending on the Category and Feature of the trial’s target. Together with the CCGP results, the generalization matrices of Task Demand corroborate that this task component engaged shared neural codes across different conditions, and that the abstract format of the variable was maintained along the trial.

Results of cross-classifying the Target Category across Task Demands and Relevant Features showed a resembling profile (Figure 4D). There was a cluster of generalization during target processing, coherent with the previous results showing that the Target Category was strongly coded in that time window (see RSA and Time-resolved MVPA results). Additional points of cross-classification during instruction encoding and between target processing and both instruction encoding and the pre-target interval were found. In accordance with what was described previously, the neural patterns of Target Category reflected an abstract format transiently during target processing, but not so evidently during the early stages of the trial. Last, the Target Relevant Feature only exhibited segregated points of significant classification that did not depict any recognizable generalization pattern (Figure 4E). As we described with the CCGP results, this task variable was only explicitly coded in the data for a very brief time period during the encoding of the third instruction screen.

### The dimensionality of the neural codes shifts along the trial

We examined the complexity of information build-up with instruction encoding, preparation and implementation across the whole trial with a decoding-based dimensionality analysis.

The results in Figure 5B show the number of dichotomies decoded significantly above chance per time-point (Rigotti et al., 2013; Bernardi et al., 2020). First, during instruction encoding, matching the onset of the second instruction’s screen until the onset of the fourth, the overall separability of the neural codes increased. We observed similar results on a second instance, concurring with target onset, and lasting for 1000 ms with a slow decrease in dimensionality towards trial completion. During the pre-target interval, in between these two time periods of higher separability, the number of dichotomies decoded decreased drastically, going back to the initial low-dimensional state.

The complementary Shattering Dimensionality index shows similar temporal dynamics. The results show an increase in SD decoding performance during instruction encoding, which diminished during the pre-target interval, and had a later peak around target onset (Figure 5C).

### The neural representational format suggests a parallel geometry favoring generalization

Building on previous research reporting representational strength of task demands and contexts in the coding of instructions (Sobrado et al., 2022; Muhle-Karbe et al., 2017), we further evaluated the spatial configuration of the activity patterns coding the two Task Demands. For that purpose, we employed model-based RSA (Kriegeskorte, 2008) and compared two alternative theoretical geometries: one where the low-dimensional structure of integration and selection was organized in parallel planes (offset model; Figure 6A left), favoring generalization of information, and another conformed by planes laying at 90 degrees (orthogonal model; Figure 6A right), reducing interference of conflicting information.

The results indicated that the offset model explained the most variance for the majority of the trial (Figure 6D). This model started predicting the pairwise distances of the conditions very early on, for a brief period during the encoding of the first instruction’s screen. Later, it had significant t-values again during the encoding of the second and third screens (for a 1300 ms window). During late encoding of the fourth screen, there was a slight increment of the t-values, with the significantly above-zero values maintained for the remainder of the trial. Prior to the presentation of the target, the results of the offset geometry exhibited a ramping-up pattern, peaking on target processing (approx. at 6000 ms).

The orthogonal model, with integration and selection laying at a 90° angle, had significant results only after target presentation. Looking at the temporal profile of its t-values, the orthogonal model started with values near chance, then the model’s explained variance diminished, with t-values below zero during the preparatory stage, instruction encoding and pre-target interval. These negative t-values hint that the orthogonal model may be negatively correlated to the neural data. Considering that offset and orthogonal models show opposing elements, these results could further suggest that neural similarity was organized by a structure more in line with a parallel geometry. After target onset, there was an increase in the model predictability, with t-values reaching significant above-zero rates. In summary, these findings suggest a geometry in the neural space in which integration and selection are represented separately, but with a configuration that allows the generalization of task information, with a shift towards orthogonality after target presentation.

## 4. Discussion

This study investigated how multi-component novel verbal instructions organize neural patterns during task preparation and execution, in a temporally-resolved manner. By characterizing the temporal unfolding of the content and geometry reflected in activity patterns, our findings demonstrate how neural codes shift in abstraction and dimensionality, corresponding to the varying nature in which the human brain encodes task-relevant components.

Our study is the first to characterize in time the neural coding of information underlying multi-component novel verbal instructions, since previous investigations primarily used fMRI (Palenciano et al., 2019a; Muhle-Karbe et al., 2017; González-García et al., 2017). Time-resolved MVPA results revealed that neural patterns carried task information during both proactive preparation and reactive control implementation (Braver et al., 2007). We observed content-specific activity patterns during instruction encoding and preparation, consistent with previous evidence of preparatory activity before target presentation (Palenciano et al., 2019a; Muhle-Karbe et al., 2017), related to the active maintenance of goal-relevant information. Specific activity patterns also appeared after target presentation, associated with reactive control processes (Palenciano et al., 2019b; González-García et al., 2017). Results of the Temporal Generalization Analysis challenged the independence of these two control modes, as we observed significant transfer of neural codes across task preparation and implementation. Two of the three variables manipulated, task demands and target animacy, showed stable underlying patterns that transferred through control epochs (Palenciano et al., 2019b), stressing the intermixed nature of neural coding for proactive and reactive control modes.

The temporal profiles of neural codes underlying different instruction components varied notably. MVPA revealed that the broader, higher-level variable of task demands was encoded earlier and maintained for longer, whereas more specific components (target category and relevant feature) did so fleetingly. This divergence was also evident in the representational space, as task demand accounted for more unique variability in shared variance analysis, over extended periods of time. Previous research has proposed hierarchical processing of task sets (Badre & Nee, 2018; Cole et al., 2011), with stronger representations for higher-level variables (Woolgar et al., 2011b; Cellier et al., 2022). Manipulating broader task demands or contexts induces highly separable representational schemes across both repetitive (Stokes et al., 2013) and novel tasks (González-García et al., 2017; Palenciano et al., 2019a). Our results extend these findings concerning lower-level variables that increased the complexity of the resulting task set: Target Category and Relevant Feature. Although category was non-essential for accurate responses in most trials, participants still coded this information, evidenced by above-chance decoding during preparation. This aligns with past findings emphasizing category information in neural patterns, even when not required for task fulfillment (Connolly et al., 2012; Soto & Ashby, 2015), reflecting category-specific anticipatory processes (Palenciano et al., 2019; Ritz et al., 2024). Part of these effects may also support preparatory mismatch detection for infrequent, unpredictable catch trials. In contrast, the temporal coding of relevant features likely engaged more intricate cognitive control processes, often linked to lower decoding accuracies (Bhandari et al., 2018), as feature conditions did not differ perceptually during target processing (targets contained both color and shape). The feature component, tied to stimulus-response mappings, potentially reflects a later step in task set hierarchy. Its less sustained coding before target presentation further supports this interpretation.

We further characterized control representations by investigating their neural geometry through abstraction and dimensionality. Cross-Condition Generalization Performance revealed that all instruction components were eventually coded in an abstract format, generalizing across different training and testing conditions. Generally, when the content of each instruction component was decoded, there was evidence of abstract coding. Thus, lower-level variables were portrayed abstractly during critical time intervals. Previous fMRI studies reported sustained abstract task representations during planning (Vaidya et al., 2021) and execution (Bhandari et al., 2024), with similar results from Deep Neural Networks (Riveland & Pouget, 2024). Among our variables, task demand exhibited more stable CCGP, corroborated by cross-classified Temporal Generalization analysis and similar fMRI results (Palenciano et al., in preparation). These findings contrast with studies lacking generalization across task contexts (e.g., Sobrado et al., 2022), consistent with conjunctive coding of information (Kikumoto & Mayr, 2020). This suggests that the generalizability of neural codes depends on computational demands, a hypothesis for future testing.

To address the specific geometry underlying our task constraints we employed RSA, establishing clear predictions on the shape of low-dimensional manifolds (simplified to three dimensions). Given the combinatorial nature of the instructions, we hypothesized that the neural coding of task demands could facilitate information sharing through a parallel geometry, rather than minimizing overlap with an orthogonal model. Our results suggest that neural space was broadly organized by an offset, parallel geometry, with integration and selection operating in separate, parallel planes. Recent evidence has reported that aligning task contexts in the neural state space promotes the generalization of conceptual information (Sheahan et al., 2021). Interestingly, we found evidence of orthogonal configuration after target presentation, which may reduce interference among competing responses (Flesch et al., 2022). Thus, while verbal instructions induced a neural format that facilitated transfer across task demands, this configuration remained somewhat constrained during implementation. A similar phenomenon of orthogonalization has been identified in the transition from evaluation to action selection (Kaufman et al., 2014; Yoo & Hayden, 2020). Orthogonalization might enable the brain to perform separate yet linked computations simultaneously (Ritz & Shenhav, 2024), reducing the risk of premature or incorrect responses (Kaufman et al., 2014). Moreover, our task manipulations generated a substantial pool of possible target combinations, and the shift towards orthogonality could enhance the encoding of distinct target attributes (Bondanelli et al., 2021). Nevertheless, our study was not designed to address these complex effects. Further research should investigate this context-dependent organization and elucidate its theoretical implications.

Regarding the temporal dynamics of neural dimensionality (Rigotti et al., 2013), we identified two key intervals characterized by more decoded dichotomies and increased Shattering Dimensionality. Studies have shown that the number of decoded dichotomies grows exponentially with the underlying dimensions of neural patterns (Rigotti et al., 2013), linked to non-linear, high-dimensional mixed selectivity (Rigotti et al., 2013; Badre et al., 2021). We observed varying dimensionality throughout the task. Once information was instructed, neural patterns became less differentiated, suggesting a compression of dimensionality that may facilitate task-relevant generalization. Nonetheless, the concurring reduction in decoding performance suggests that information was not explicitly maintained during this interval. Upon target presentation, activity transitioned back to a high-dimensional state, potentially reducing interference among target combinations. Studies employing repetitive tasks show rapid dimensionality expansion after stimuli presentation, with neural codes morphing to higher-dimensional states (Parthasarathy et al., 2017; Aoi et al., 2020) linked to improved behavioral efficiency (Kikumoto et al., 2024a). Our results highlight how information complexity shifts dynamically across instruction encoding and implementation stages, with an expansion-compression interplay that may serve as a control mechanism to solve the “curse of dimensionality” (Bellman, 1957) while minimizing interference.

The overall pattern of dimensionality results contributes to the debate on the neural geometry of novel tasks. Some authors argue that dimensionality reduction occurs over learning (Farrell et al., 2022; Wójcik et al., 2023), while others suggest that novel settings start with fewer dimensions and abstract codes (Cole et al., 2013; Verbeke & Verguts, 2022). In contrast, we observed an interesting pattern featuring high dimensionality alongside generalization across conditions. Discrepancies may stem from the nature of the tasks used, as most previous dimensionality studies involve slower trial-and-error learning (Wójcik et al., 2023; Flesch et al., 2022), whereas verbally instructed tasks create complex task sets early on. Additionally, task settings with constant structure or high complexity tend to favor generalization (Aoi et al., 2020; Woolgar et al., 2015), both of which were present in the current experiment. Our results do not support an initial high (Wójcik et al., 2023) or low-dimensional state (Verbeke & Verguts, 2022), but instead reveal a more complex representational landscape for novel recombination scenarios.

Despite its contributions, our study is not without limitations. Adapting the paradigm to EEG required splitting instructions into sequential words, while most previous studies present instructions all at once (González-García, 2017; Palenciano et al., 2019a). This sequential presentation may have affected the structure underlying the neural codes observed, restricting comparisons with previous results. Regarding the novelty in our paradigm, although the whole instructions were different and new on each trial, the overarching task structure remained consistent across the session, and elements were recursively reused, as commonly done in such studies (Cole et al., 2011; Ruge et al., 2019). Thus, practice might have played a role. Ruge et al. (2019) found significant changes in task representations from early to late trials, and decreases in representational strength due to familiarity have been reported (Woolgar et al., 2011a) accompanied with a shift toward stronger high-level task conjunctions (Kikumoto et al., 2024b). Unfortunately, however, our study lacked enough trials to explore practice using multivariate techniques. Nonetheless, our approach focused not only on how novel task sets are configured in single trials, but primarily on how the brain compositionally adds sub-components to build complex task sets over time. Finally, EEG limits conclusions about the spatial sources of brain activation. Fusion analysis of EEG and fMRI data (Cichy & Oliva, 2020) could help pinpoint the regions involved in these temporal dynamics.

In conclusion, this study sheds light on how information is represented and flexibly reconfigured to meet demands instructed verbally. Results show that re-used instruction content is coded in an abstract format, enabling generalization across contexts, while using high-dimensional patterns that vary with stages of information processing. Further studies could extrapolate these findings to novel scenarios with an overall different task structure and complement them with more spatially precise neuroimaging techniques.

## CRediT author contribution

PP: Conceptualization, Methodology, Software, Formal Analysis, Investigation, Data Curation, Writing - Original Draft Preparation, Writing - Review & Editing, Visualization. AFP: Conceptualization, Methodology, Software, Investigation, Resources, Data Curation, Writing - Review & Editing, Supervision. CGG: Conceptualization, Writing - Review & Editing, Supervision. MR: Conceptualization, Resources, Writing - Review & Editing, Supervision, Project administration, Funding acquisition.

## Data and code availability

Raw behavioral and EEG data will be made available upon publication. Preprocessing and analysis codes can be found at https://github.com/pena-p/inst-comp-eeg.git

## Acknowledgments

This research was supported by grant PID2022-138940NB-100 awarded to MR and grant PID2023-151911NA-I00 awarded to AFP, funded by MCIN/EI/10.13039/501100011033/ and FEDER, UE. AFP was supported by Grant PAIDI21_00207 of the Andalusian Autonomic Government. CCGP was supported by Project PID2020-116342GA-I00 funded by MCIN/AEI/10.13039/501100011033, and Grant RYC2021-033536-I funded by MCIN/AEI/10.13039/501100011033 and by the European Union NextGeneration EU/PRTR. The Mind, Brain and Behavior Research Center receives funding from grants CEX2023-001312-M by MCIN/AEI /10.13039/501100011033 and UCE-PP2023-11 by the University of Granada. We are grateful to Yulya Bedritskaya and Marta Becerra Losada for assistance with data collection, and Chiara Avancini for assistance with the EEG equipment setup.

