## Supplementary_Material for "Novel verbal instructions recruit abstract neural patterns of time-variable information dimensionality"

| Nº | TRAINING SET | | | | | | | TESTING SET | | |
|---|---|---|---|---|---|---|---|---|---|---|
| | Class A | | | | Class B | | | Class A | | Class B |
| 1 | 2 | 3 | 4 | vs. | 2 | 3 | 4 | 1 | vs. | 1 |
| 2 | 2 | 3 | 4 | vs. | 1 | 3 | 4 | 1 | vs. | 2 |
| 3 | 2 | 3 | 4 | vs. | 1 | 2 | 4 | 1 | vs. | 3 |
| 4 | 2 | 3 | 4 | vs. | 1 | 2 | 3 | 1 | vs. | 4 |
| 5 | 1 | 3 | 4 | vs. | 2 | 3 | 4 | 2 | vs. | 1 |
| 6 | 1 | 3 | 4 | vs. | 1 | 3 | 4 | 2 | vs. | 2 |
| 7 | 1 | 3 | 4 | vs. | 1 | 2 | 4 | 2 | vs. | 3 |
| 8 | 1 | 3 | 4 | vs. | 1 | 2 | 3 | 2 | vs. | 4 |
| 9 | 1 | 2 | 4 | vs. | 2 | 3 | 4 | 3 | vs. | 1 |
| 10 | 1 | 2 | 4 | vs. | 1 | 3 | 4 | 3 | vs. | 2 |
| 11 | 1 | 2 | 4 | vs. | 1 | 2 | 4 | 3 | vs. | 3 |
| 12 | 1 | 2 | 4 | vs. | 1 | 2 | 3 | 3 | vs. | 4 |
| 13 | 1 | 2 | 3 | vs. | 2 | 3 | 4 | 4 | vs. | 1 |
| 14 | 1 | 2 | 3 | vs. | 1 | 3 | 4 | 4 | vs. | 2 |
| 15 | 1 | 2 | 3 | vs. | 1 | 2 | 4 | 4 | vs. | 3 |
| 16 | 1 | 2 | 3 | vs. | 1 | 2 | 3 | 4 | vs. | 4 |

**Figure 4-1.** Split of the data into training and test sets for the CCGP analysis. The training set consisted of 3 conditions per side of the classification, and the test set of 1 condition per side of the classification. In total, after all unique test combinations were implemented, 16 possible groupings of the data were used.

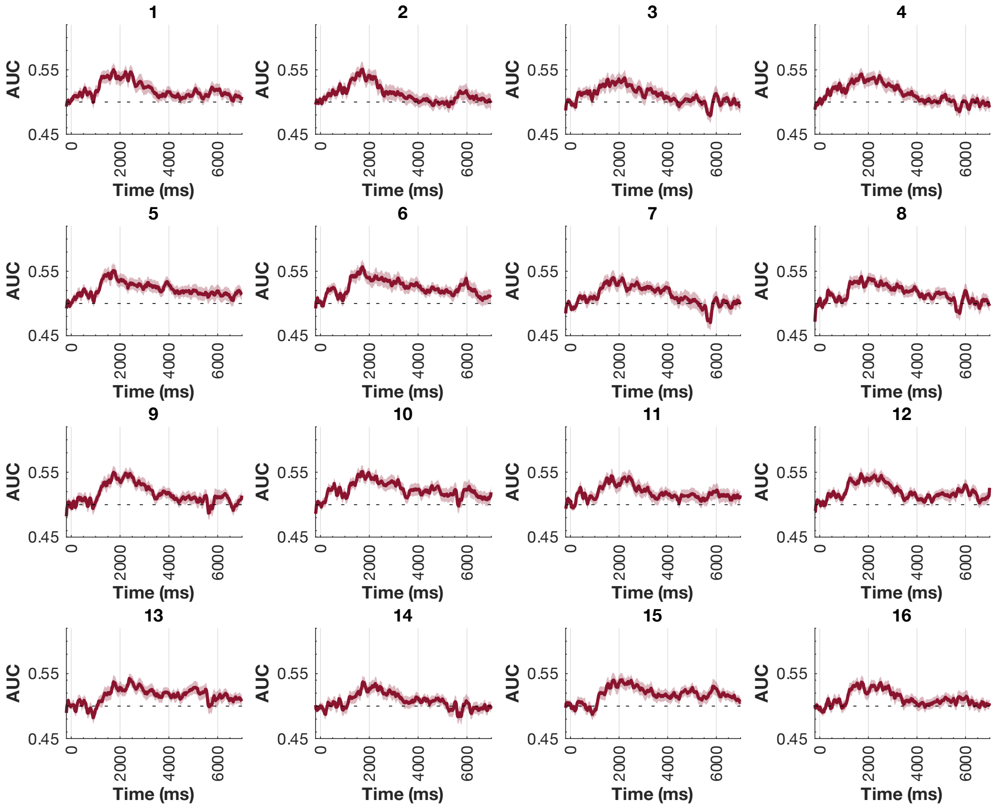


**Figure 4-2.** Cross-classification results of Task Demand per combination of train and test sets for Cross-Classification Generalization Performance (CCGP) Analysis. Each of the 16 combinations has its own subplot of cross-decoding performance (indicated by the number on top). The darker magenta lines represent the Area Under the Curve (AUC) over time (in ms).

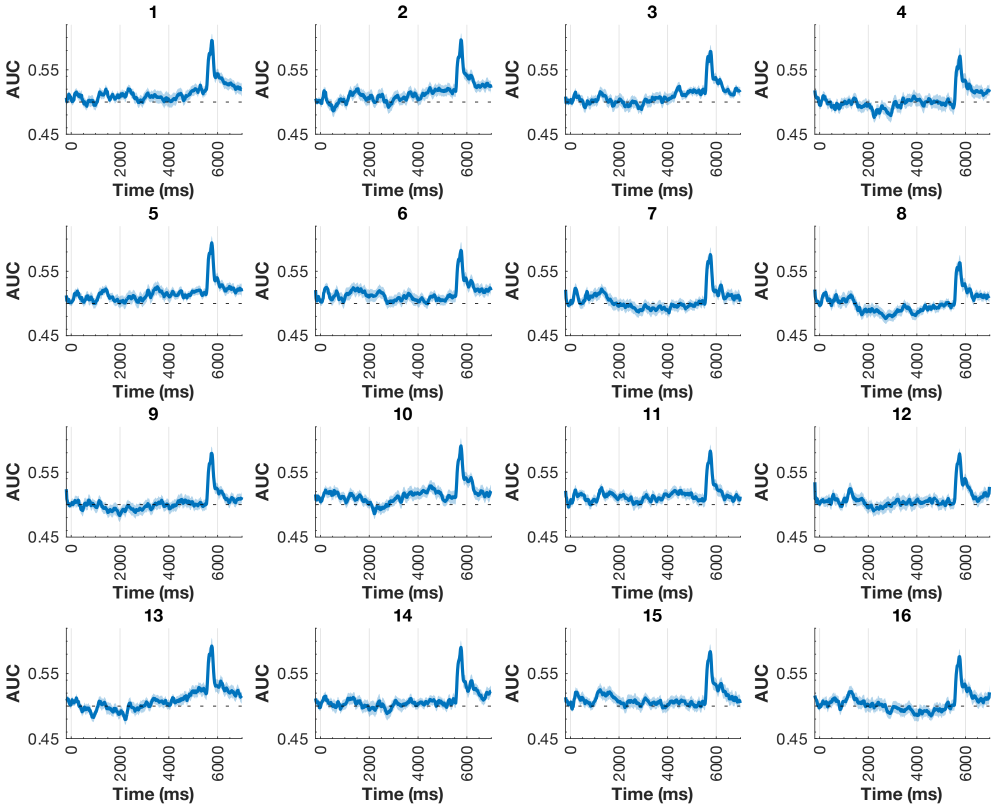


**Figure 4-3.** Cross-classification results of Target Category per combination of train and test sets for Cross-Classification Generalization Performance (CCGP) Analysis. Each of the 16 combinations has its own subplot of cross-decoding performance (indicated by the number on top). The darker blue lines represent the Area Under the Curve (AUC) over time (in ms).

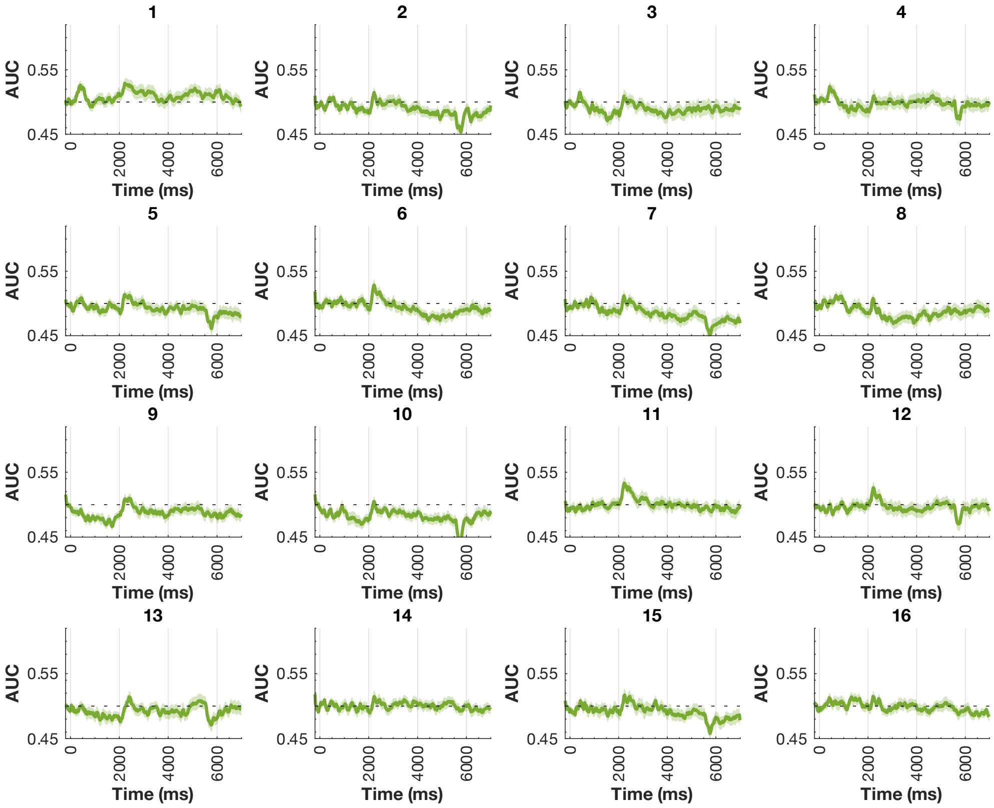


**Figure 4-4.** Cross-classification results of Target Relevant Feature per combination of train and test sets for Cross-Classification Generalization Performance (CCGP) Analysis. Each of the 16 combinations has its own subplot of cross-decoding performance (indicated by the number on top). The darker green lines represent the Area Under the Curve (AUC) over time (in ms).

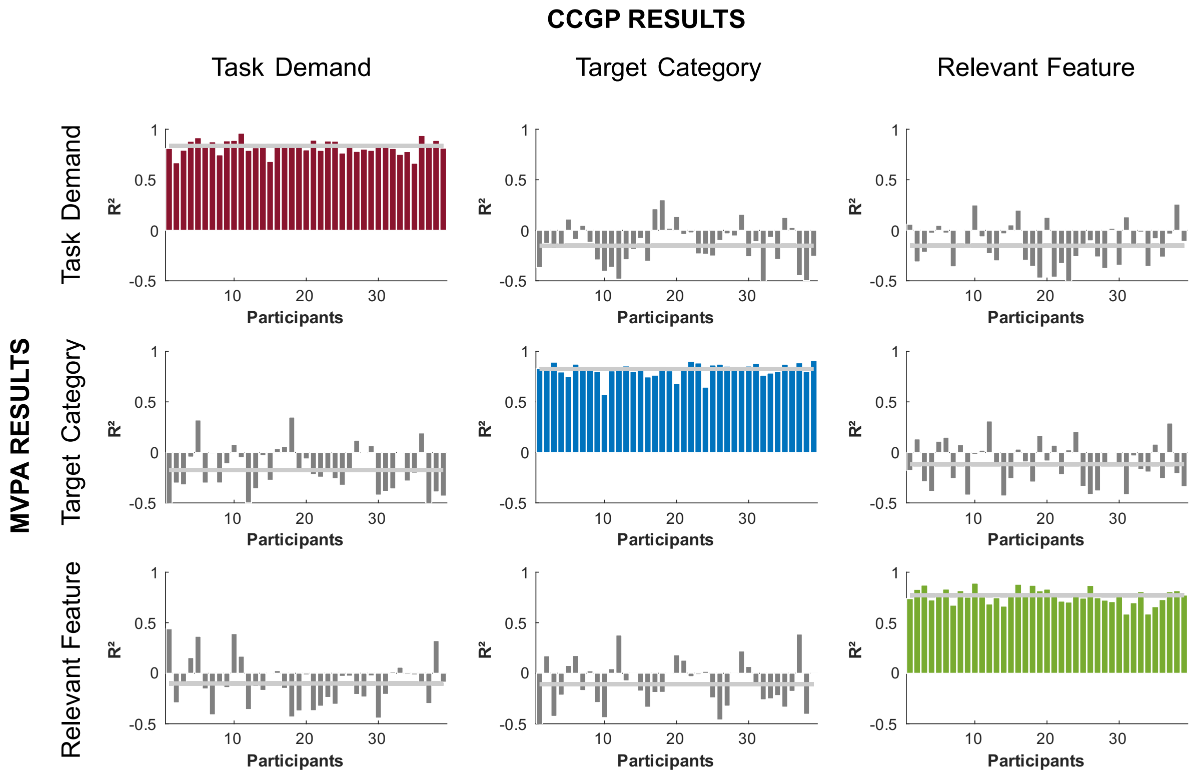


**Figure 4-5.** Correlation of regular MVPA and CCGP results. Histograms of Pearson coefficients across participants obtained by comparing the three variables pair-wise. Each row of histograms refers to the results of regular MVPA of each of the three variables (row 1: Task Demand, row 2: Target Category, row 3: Target Relevant Feature). Each column of the histograms refers to the results of CCGP of each of the three variables (column 1: Task Demand, column 2: Target Category, column 3: Target Relevant Feature). Each bar corresponds to the Pearson coefficient of a participant in a specific pair-wise comparison, the grey horizontal line represents the averaged Pearson coefficient of that comparison after performing Fisher's method (Fisher, 1970). The diagonal histograms correspond to the correlation of regular MVPA and CCGP results of the same variable: Task Demand (magenta), Target Category (blue), and Target Relevant Feature (green). The remaining histograms result from comparing different variables (i.e., MVPA of Task Demand and CCGP of Target Category), and were used as a control to ensure that the diagonal correlations were not due to the structure of the noise.

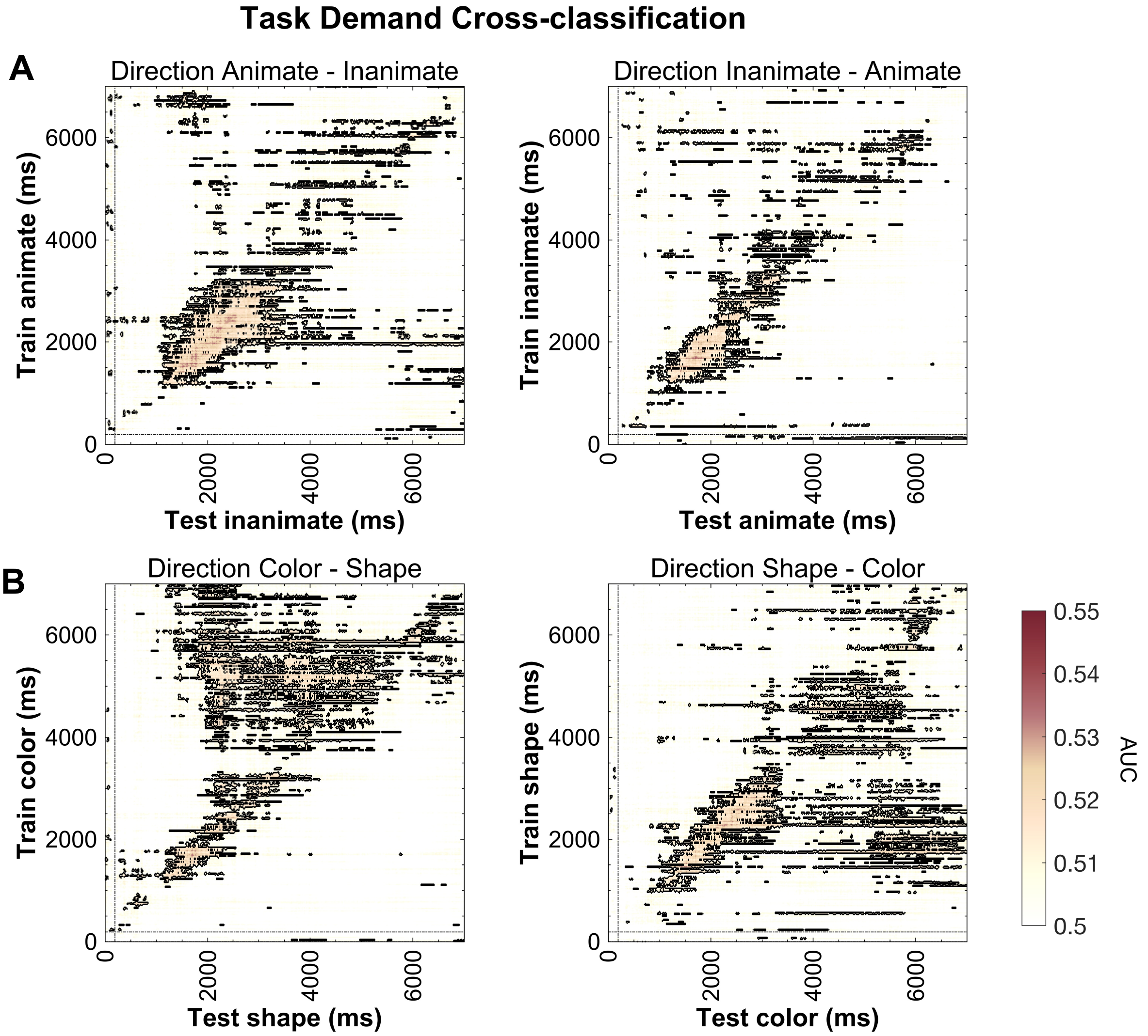


**Figure 4-6.** Cross-classification temporal generalization results per direction of Task Demand across different contexts. **(A)** Cross-classification of integration vs. selection across different Target Categories, when trained on animate and tested on inanimate (left) and when trained on inanimate and tested on animate (right). **(B)** Cross-classification integration vs. selection across different Target Relevant Features, when trained on color and tested on shape (left) and when trained on shape and tested on color (right). The vertical axis indicates the training time bins, and the horizontal axis the testing time bins. The color bar indicates the AUC value of the classifier, black lines denote significant clusters.

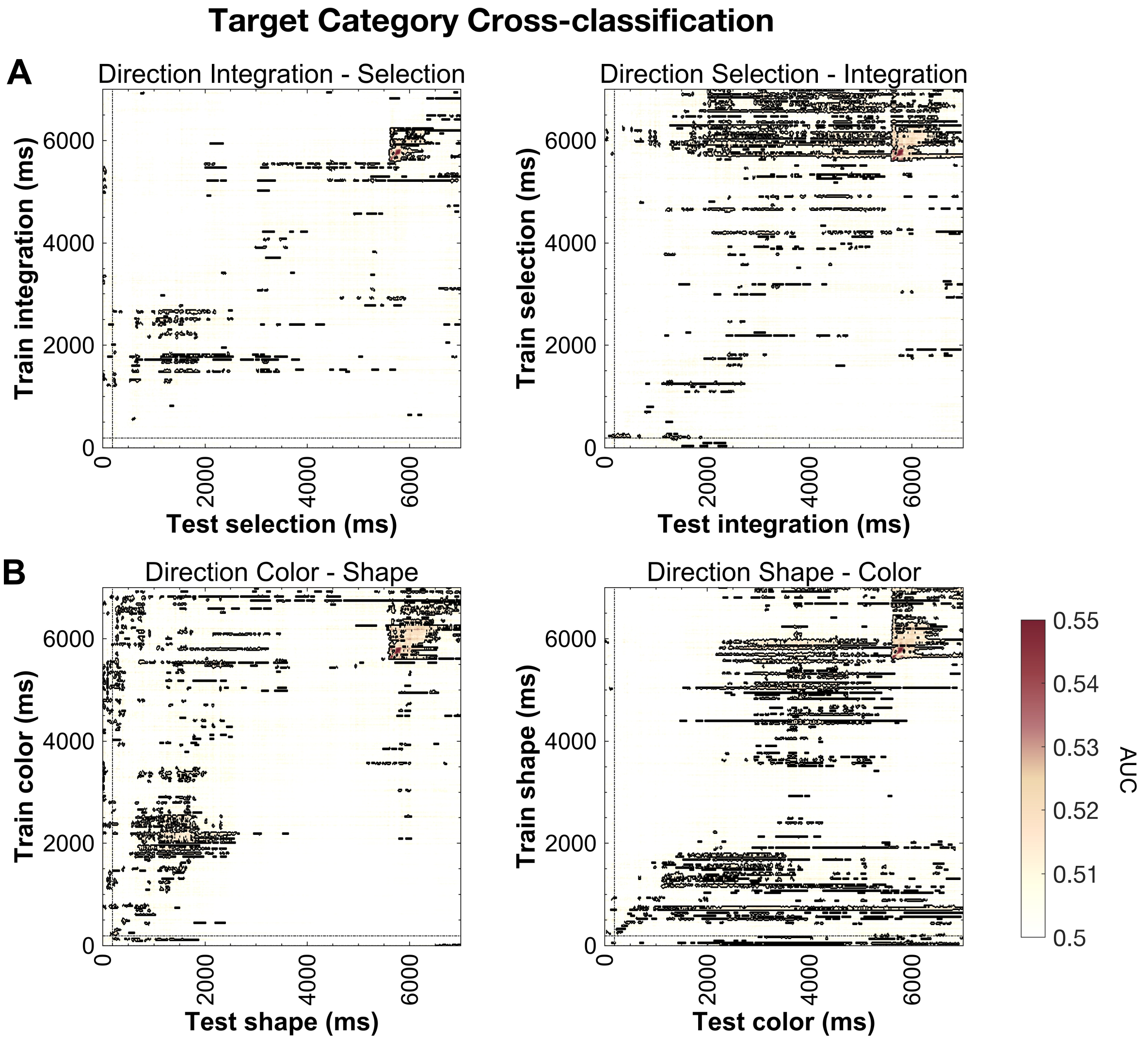


**Figure 4-7.** Cross-classification temporal generalization results per direction of Target Category across different contexts. **(A)** Cross-classification of animate vs. inanimate across different Task Demands, when trained on integration and tested on selection (left) and when trained on selection and tested on integration (right). **(B)** Cross-classification of animate vs. inanimate across different Target Relevant Features, when trained on color and tested on shape (left) and when trained on shape and tested on color (right). The vertical axis indicates the training time bins, and the horizontal axis the testing time bins. The color bar indicates the AUC value of the classifier, black lines denote significant clusters.

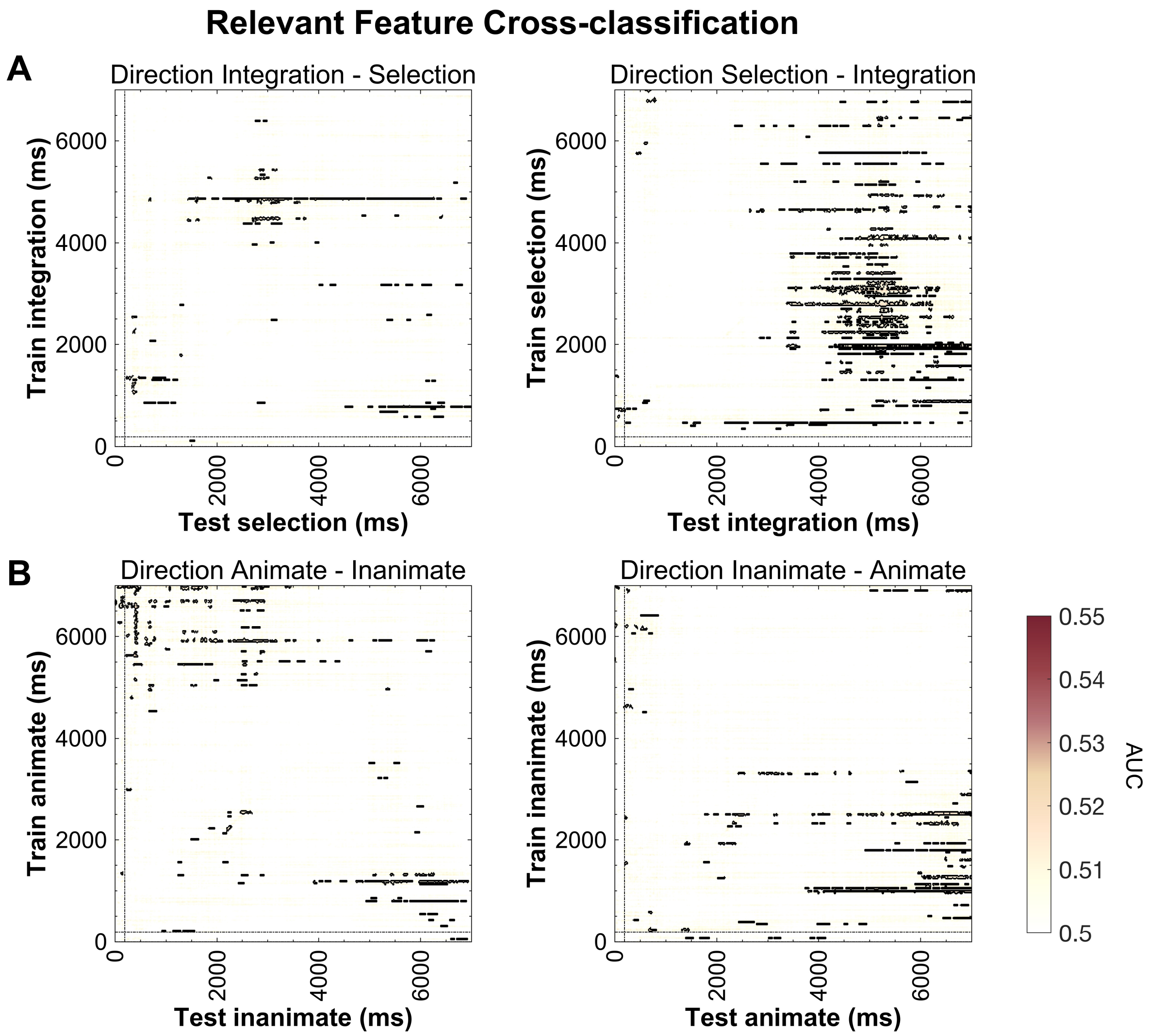


**Figure 4-8.** Cross-classification temporal generalization results per direction of Target Relevant Feature across different contexts. **(A)** Cross-classification of color vs. shape across different Task Demands, when trained on integration and tested on selection (left) and when trained on selection and tested on integration (right). **(B)** Cross-classification of color vs. shape across different Target Categories, when trained on animate and tested on inanimate (left) and when trained on inanimate and tested on animate (right). The vertical axis indicates the training time bins, and the horizontal axis the testing time bins. The color bar indicates the AUC value of the classifier, black lines denote significant clusters.

| | CLASSIFICATION DISPOSITION | | | | | | | | | |
|---|---|---|---|---|---|---|---|---|---|---|
| Nº | Side A of Dichot. | | | | | Side B of Dichot. | | | | Variable Name |
| 1 | 1 | 2 | 3 | 4 | vs. | 5 | 6 | 7 | 8 | Task Demand |
| 2 | 1 | 2 | 3 | 5 | vs. | 4 | 6 | 7 | 8 | N/A |
| 3 | 1 | 2 | 3 | 6 | vs. | 4 | 5 | 7 | 8 | N/A |
| 4 | 1 | 2 | 3 | 7 | vs. | 4 | 5 | 6 | 8 | N/A |
| 5 | 1 | 2 | 3 | 8 | vs. | 4 | 5 | 6 | 7 | N/A |
| 6 | 1 | 2 | 4 | 5 | vs. | 3 | 6 | 7 | 8 | N/A |
| 7 | 1 | 2 | 4 | 6 | vs. | 3 | 5 | 7 | 8 | N/A |
| 8 | 1 | 2 | 4 | 7 | vs. | 3 | 5 | 6 | 8 | N/A |
| 9 | 1 | 2 | 4 | 8 | vs. | 3 | 5 | 6 | 7 | N/A |
| 10 | 1 | 2 | 5 | 6 | vs. | 3 | 4 | 7 | 8 | Target Category |
| 11 | 1 | 2 | 5 | 7 | vs. | 3 | 4 | 6 | 8 | N/A |
| 12 | 1 | 2 | 5 | 8 | vs. | 3 | 4 | 6 | 7 | N/A |
| 13 | 1 | 2 | 6 | 7 | vs. | 3 | 4 | 5 | 8 | N/A |
| 14 | 1 | 2 | 6 | 8 | vs. | 3 | 4 | 5 | 7 | N/A |
| 15 | 1 | 2 | 7 | 8 | vs. | 3 | 4 | 5 | 6 | N/A |
| 16 | 1 | 3 | 4 | 5 | vs. | 2 | 6 | 7 | 8 | N/A |
| 17 | 1 | 3 | 4 | 6 | vs. | 2 | 5 | 7 | 8 | N/A |
| 18 | 1 | 3 | 4 | 7 | vs. | 2 | 5 | 6 | 8 | N/A |
| 19 | 1 | 3 | 4 | 8 | vs. | 2 | 5 | 6 | 7 | N/A |
| 20 | 1 | 3 | 5 | 6 | vs. | 2 | 4 | 7 | 8 | N/A |
| 21 | 1 | 3 | 5 | 7 | vs. | 2 | 4 | 6 | 8 | Relevant Feature |
| 22 | 1 | 3 | 5 | 8 | vs. | 2 | 4 | 6 | 7 | N/A |
| 23 | 1 | 3 | 6 | 7 | vs. | 2 | 4 | 5 | 8 | N/A |
| 24 | 1 | 3 | 6 | 8 | vs. | 2 | 4 | 5 | 7 | N/A |
| 25 | 1 | 3 | 7 | 8 | vs. | 2 | 4 | 5 | 6 | N/A |
| 26 | 1 | 4 | 5 | 6 | vs. | 2 | 3 | 7 | 8 | N/A |
| 27 | 1 | 4 | 5 | 7 | vs. | 2 | 3 | 6 | 8 | N/A |
| 28 | 1 | 4 | 5 | 8 | vs. | 2 | 3 | 6 | 7 | N/A |
| 29 | 1 | 4 | 6 | 7 | vs. | 2 | 3 | 5 | 8 | N/A |
| 30 | 1 | 4 | 6 | 8 | vs. | 2 | 3 | 5 | 7 | N/A |
| 31 | 1 | 4 | 7 | 8 | vs. | 2 | 3 | 5 | 6 | N/A |
| 32 | 1 | 5 | 6 | 7 | vs. | 2 | 3 | 4 | 8 | N/A |
| 33 | 1 | 5 | 6 | 8 | vs. | 2 | 3 | 4 | 7 | N/A |
| 34 | 1 | 5 | 7 | 8 | vs. | 2 | 3 | 4 | 6 | N/A |
| 35 | 1 | 6 | 7 | 8 | vs. | 2 | 3 | 4 | 5 | N/A |

**Figure 5-1.** Splitting of the data into balanced dichotomies (4 vs. 4) for dimensionality analysis. A total of 35 dichotomies are obtained after all possible combinations of the 8 conditions are illustrated.

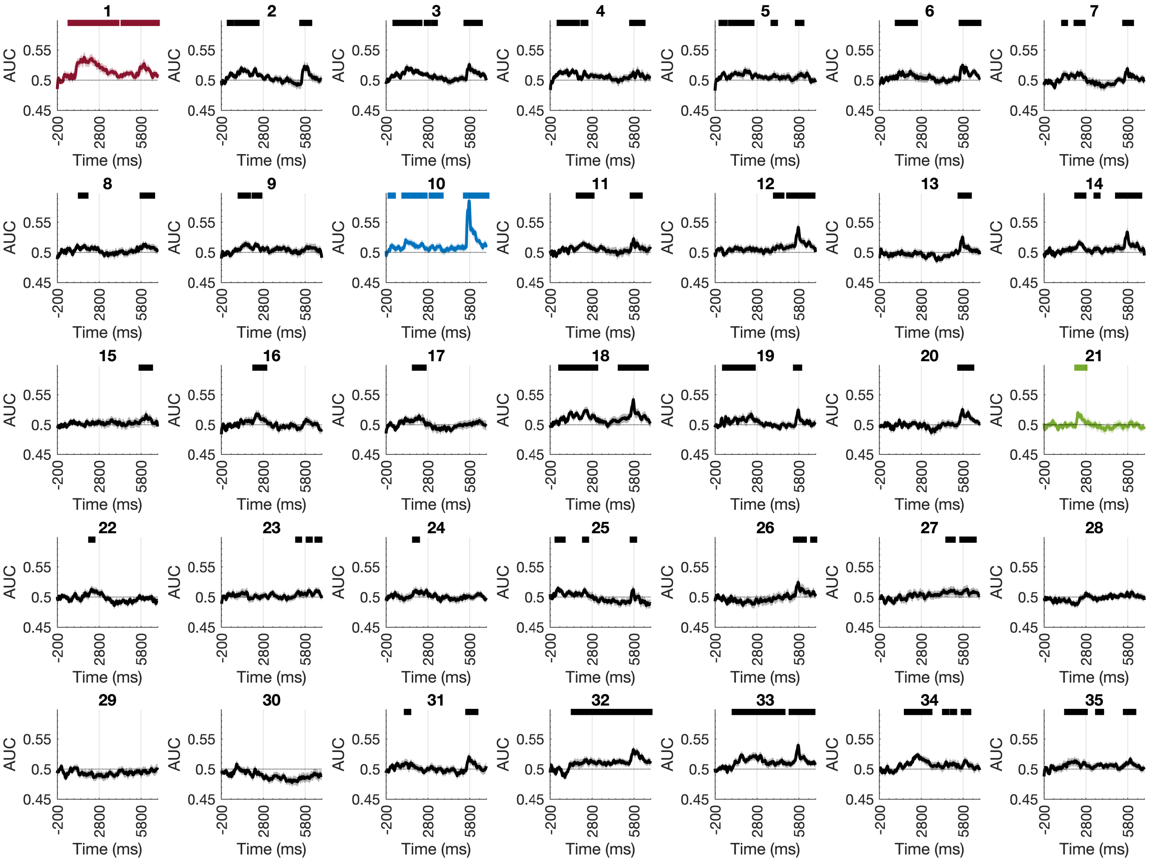


**Figure 5-2.** Decoding results of Shattering Dimensionality per dichotomy of the conditions. Each of the 35 dichotomies has its own subplot of decoding performance (indicated by the number on top). The darker lines represent the Area Under the Curve (AUC) over time (in ms), the colored dichotomies refer to the splits of data that correspond to our three main task variables (magenta: Task Demand, blue: Target Category, and green: Target Relevant Feature). Lighter shading refers to the standard error associated with the mean of the performance measure. The horizontal-colored lines refer to the statistical significance against chance (dotted line) after implementing cluster-based permutation analysis per dichotomy (necessary to obtain the number of significantly decoded dichotomies, Figure 6C).
